# Virion-associated US28 rapidly modulates Akt activity to suppress HCMV lytic replication in monocytes

**DOI:** 10.1101/2023.09.05.556359

**Authors:** Jamil Mahmud, Brittany W. Geiler, Juthi Biswas, Michael J. Miller, Julia E. Myers, Stephen M. Matthews, Amanda B. Wass, Christine M. O’Connor, Gary C. Chan

**Author notes:** Correspondence and inquiries regarding the manuscript should be addressed to: Gary C. Chan, Ph.D., Department of Microbiology and Immunology, SUNY Upstate Medical University, 750 East Adams Street, Syracuse, NY 13210.

## Abstract

Establishing a non-productive quiescent/silent infection within monocytes is essential for spread of human cytomegalovirus (HCMV). Yet, how HCMV establishes a quiescent infection in monocytes remains unclear. US28 is a viral G protein-coupled receptor (GPCR) essential for silent infections within cells of the myeloid lineage. We found virion-associated US28 was rapidly delivered to monocytes, while *de novo* synthesized US28 was delayed for several days. A recombinant mutant virus lacking US28 (US28Δ) was unable to establish a quiescent infection, resulting in a fully productive lytic replication cycle. Mechanistically, viral entry of US28Δ phosphorylated Akt at both serine 473 (S473) and threonine 308 (T308), which contrasted with the site-specific phosphorylation of Akt at S473 following WT infection. Preventing Akt bi-phosphorylation prevented lytic replication of US28Δ, and ectopic expression of a constitutively phosphorylated Akt variant triggered lytic replication of WT infection. Our data demonstrate that virion-delivered US28 fine-tunes Akt activity to permit HCMV infection to enter a quiescent state following primary infection of monocytes.

## Introduction

Infection with human cytomegalovirus (HCMV) is highly prevalent, with seropositivity reaching 80% in developed countries and upwards of 100% in developing countries (*1*). HCMV infection results in lifelong latent infection, with more than 50% individuals aged 6 years harboring HCMV, which rises to 90% by the age of 75 in the United States (*2, 3*). In healthy individuals, primary HCMV infection is generally asymptomatic, and latency is tightly controlled by the immune system. However, HCMV primary infection or reactivation is associated with significant morbidity and mortality in the immunonaïve, such as neonates, and the immunocompromised, such as HIV patients and transplant recipients (*4–7*). HCMV-associated inflammatory diseases within these patients are often widespread and can lead to multi-organ failure.

The myeloid compartment is central to the HCMV viral dissemination and persistence strategy. Following a primary infection, HCMV replicates in oral epithelial cells and spreads to peripheral blood monocytes where the virus establishes a quiescent infection, which is defined by 1) the suppression of virus replication and 2) a spontaneously reactivation of the virus after 2-3 weeks of infection in the absence of external stimuli (*8, 9*). Infection directly stimulates monocytes to travel to distal end-organ sites and differentiate into long-lived macrophages permissive for viral replication. Infected monocytes that journey to the bone marrow ultimately transfer infection to CD34+ hematopoietic progenitor cells (HPCs) in order to establish a lifelong reservoir of latency. Based on this model of viral dissemination, primary infected monocytes are a critical lynchpin to viral dissemination by linking initial lytic infection to life-long latent infection in CD34+ cells. Suppression of initial lytic infection in primary infected monocytes is essential for the immune evasion and spread of HCMV within the host, as viral mutants unable to establish latency or wild type HCMV forced out of latency can be targeted by the host immune response (*10, 11*). Yet, little is known about the mechanisms underpinning the establishment of a quiescent infection within monocytes.

HCMV G protein-coupled receptor (GPCR) US28 is found within mature virions (*12*) and expressed during latent (*10, 12–19*) as well as lytic infection (e.g. refs (*20–22*)). US28 acts as a signaling molecule to enhance cellular proliferation, chemotactic and mitotic processes (discussed in (*23*)), and promotes cellular migration (*24–26*) through activation of different migratory factors, including Pyk2 and RhoA (*27, 28*). US28 also modulates multiple cellular pathways during lytic infection, including calcium signaling (*20, 21*), FAK/Src (*29*), PLC (*20, 30*), COX-2 (*31*), STAT3 (*32*), Akt, ERK1/2, eNOS (*33*), beta-catenin (*34*). Importantly, US28 is essential for viral latency/quiescence in cells of the myeloid lineage, including CD34+ HPCs (*12, 14, 15, 17*), THP1 cells (*10, 15, 35*), Kasumi-3 cells (*12, 14, 15*), and monocytes (*10, 36, 37*). During latency, US28-mediated signaling silences the major immediate-early promoter (MIEP) (*10, 12, 14, 15*), a key promoter in the latent-to-lytic switch that regulates the expression of immediate-early 1 (IE1) and immediate-early 2 (IE2) viral proteins, encoded by *UL123* and *UL122*, respectively. In Kasumi-3 cells and CD34+ HPCs, US28 attenuates cellular fos (c-fos) to prevent AP-1 transcription factor-dependent activation of the MIEP (*15*). Additionally, studies in THP-1 cells revealed the importance of US28-mediated downregulation of MAPK and NFκB pathways in establishing latency (*10*). Similar to latent infection, expression of viral lytic proteins, such as IE1, are inhibited during quiescent infection of monocytes (*8, 9, 38, 39*). Although virion-associated US28 is important for the establishment of latency in Kasumi-3 cells and CD34+ HPCs (*15*), the role of US28-mediated regulation of the MIEP during the establishment of a quiescent infection in monocytes remains to be elucidated.

In this study, we demonstrate that virion-associated US28 modulates Akt activity to allow for the establishment of a quiescent infection in primary, HCMV-infected monocytes. Specifically, we found that infection of peripheral blood monocytes with recombinant mutant viruses lacking US28 (US28Δ) or defective in US28-mediated signaling resulted in the rapid expression of IE, early (E), and late (L) viral proteins. Monocytes infected with US28-complemented US28Δ virus, which is able to deliver virion-associated US28 during infection while incapable of synthesizing *de novo* US28 in the infected cells, had reduced levels of *UL123*, indicating virion-associated US28 is sufficient to inhibit IE gene expression during early infection. Mechanistically, we found US28 was necessary and sufficient to limit EGFR activation induced by WT infection, which led to the site-specific phosphorylation of Akt at the S473 residue. In contrast, US28Δ-infected monocytes exhibited increased EGFR activity relative to WT infection as well as Akt phosphorylation at both S473 and T308, which is required for full Akt activity (*40, 41*). Importantly, stimulation of Akt phosphorylation at both S473 and T308 during WT infection of monocytes initiated IE1 synthesis, while suppression of either S473 or T308 phosphorylation during US28Δ infection attenuated IE1 expression. We previously showed that the partial Akt activity facilitated by S473 phosphorylation alone is required for the long-term survival of HCMV-infected monocytes (*42, 43*). We now demonstrate that preventing the full activation of Akt, via US28-mediated reduction of T308 Akt phosphorylation, is also crucial to ensuring IE protein expression in not initiated during the establishment of a quiescent infection within monocytes.

## Results

### US28-mediated signaling promotes HCMV quiescent infection within monocytes

US28 expression is essential for HCMV latency in CD34+ HPCs (*12, 14, 15, 17*), THP1 cells (*10, 15, 35*), Kasumi-3 cells (*12, 14, 15*), and monocytes (*10, 36, 37*). Accordingly, we found infection of CD14+ primary peripheral blood monocytes with an HCMV mutant lacking US28 (US28Δ) failed to establish a quiescent infection, leading to rapid IE1 protein expression at 24 through 48 hours post-infection (hpi) similar to WT infection in replication permissive fibroblasts (**Fig. 1A to C**). In addition to IE protein expression, infection of monocytes with US28Δ also resulted in early (E) and late (L) protein expression (**Fig. S1**), indicating progression through the late stages of the lytic replication cycle. To determine the importance of US28 signaling in inhibiting IE1, we used two additional US28-recombinant viral strains with altered signaling capabilities (*15*). TB40/E*mCherry*-US28ΔN-3xF (ΔN) is a chemokine binding domain-deficient mutant that lacks amino acids (aa) 2-16, thereby preventing US28’s interaction with many of its ligands (*44–46*). TB40/E*mCherry*-US28-R129A-3xF (R129A) harbors a point mutation in the ‘DRY’ motif at aa 129 to which G proteins couple, rendering R129A a G protein-coupling deficient mutant (*47, 48*). We confirmed that both ΔN and R129A mutants express recombinant US28 in the virion (**Fig. 1D**). Similar to US28Δ, infection with either signaling mutant resulted in IE1 expression in monocytes (**Fig. 1E, F**), which is consistent with previous findings in THP-1, Kasumi-3 cells, and CD34+ HPCs (*10, 15, 16*), suggesting these mutants fail to undergo quiescence in monocytes. It should be pointed out that monocytes infected with R129A and ΔN showed a higher expression of IE1 compared to US28Δ. Why IE1 expression was less robust in US28Δ-infected monocytes relative to the signaling mutants is unclear, although a possibility is that signaling deficient US28 may act as a scaffold to enhance the formation of signaling complexes during viral entry. Regardless, our data indicate both ligand binding, as well as G protein-mediated signaling via US28, are pivotal to averting IE1 expression in monocytes.

**Fig 1.**
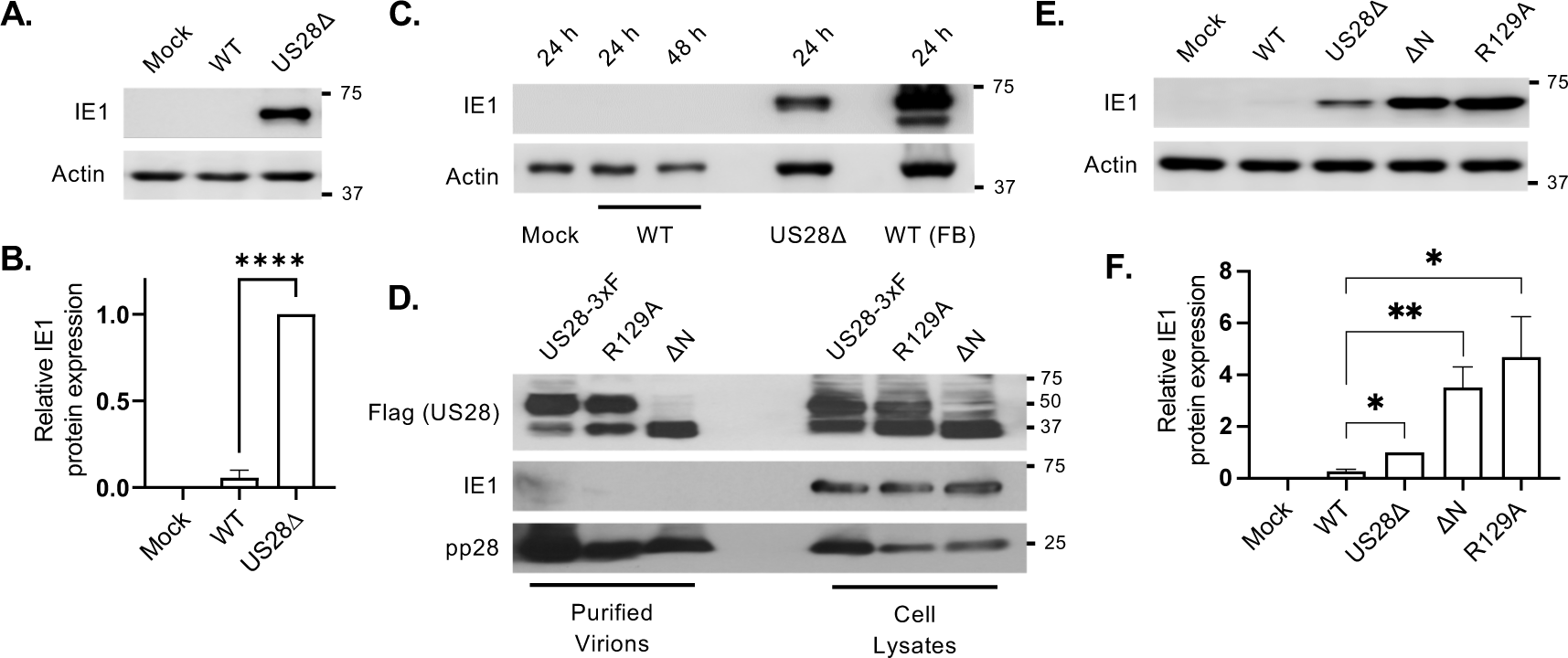
US28 mediated signaling inhibits IE1 expression in monocytes. **(A to C)** Primary human peripheral blood monocytes or primary human fibroblasts (FB) were infected (MOI = 1) with mock, wildtype (WT), or US28Δ for 24 h or 48 h. IE1 expression was detected by western blot. **(D)** NuFF-1 cells were infected (MOI= 1) with US28-3xF, R129A or ΔN. After 100% cytopathic effect was observed, virions were purified from supernatants and cell lysates collected. Incorporation of recombinant US28 into mature virions was detected by western blotting using an anti-flag antibody. IE1 and pp28 staining were used as controls. **(E)** Monocytes were infected (MOI = 1) with mock, wildtype (WT), US28Δ, ΔN or R129A, for 24 h. IE1 expression was detected by western blot. All membranes were probed for β-actin as a loading control. Western blots are representative of at least 3 independent experiments. Densitometry analysis was performed using Image Lab software (Bio-Rad) from at least 3 independent experiments **(F)**. Statistical significance was measured using one-way ANOVA and t-test; ****P < 0.0001, **P < 0.005, *P<0.05.

Next, we sought to investigate the kinetics of US28-mediated regulation of IE1 expression in US28Δ-infected monocytes. While IE1 expression was restricted in WT-infected monocytes, IE1 protein expression in US28Δ-infected monocytes was detectable as early as 6 hpi (**Fig. 2A**), which was similar to the IE1 expression kinetics in both WT- and US28Δ-infected fibroblasts (**Fig. 2B**). These results further confirm previous studies demonstrating US28 is dispensable for *in vitro* infection of fibroblasts (e.g., refs. (*12, 20, 21, 49, 50*)). We next ensured the absence of IE1 expression following WT HCMV infection was not due to reduced viral binding or entry into monocytes. First, we performed binding assays at 4°C and confirmed both WT and US28Δ bound monocytes to the same extent (**Fig. S2A**). Additionally, the absence of US28 had no effect on viral entry, as both WT and US28Δ equally entered monocytes (**Fig. S2B**). As a control, viral entry was inhibited by pre-treating monocytes with AG1478, an EGFR inhibitor known to block HCMV entry into monocytes at high concentrations (*51*), which reduced entry of both WT and US28Δ. Next, we confirmed that translocation of the viral genome to the nucleus was also not impaired by the deletion of US28. Flow cytometric analysis of mCherry expression, which is driven from an independent SV40 promoter inserted in an intergenic region of the HCMV genome (*52*), did not reveal any significant difference in the percentage of mCherry positive cells or mCherry fluorescence intensity between WT- and US28Δ-infected monocytes (**Fig. S2C, S2D**). As differentiated monocytes support lytic HCMV infection (reviewed in (*38*)), we next examined if loss of US28 enhances differentiation of the infected cells. Expression of several macrophage-associated cell-surface markers were similar in WT- and US28Δ-infected monocytes at early and late stages of infection **(Fig S3)**, suggesting lytic infection in US28Δ-infected cells is not due to an acceleration in the monocyte-to-macrophage differentiation process. Collectively, our results indicate US28 impedes IE1 expression upon entry into monocytes in a ligand- and G protein-coupling-dependent fashion.

**Fig 2.**
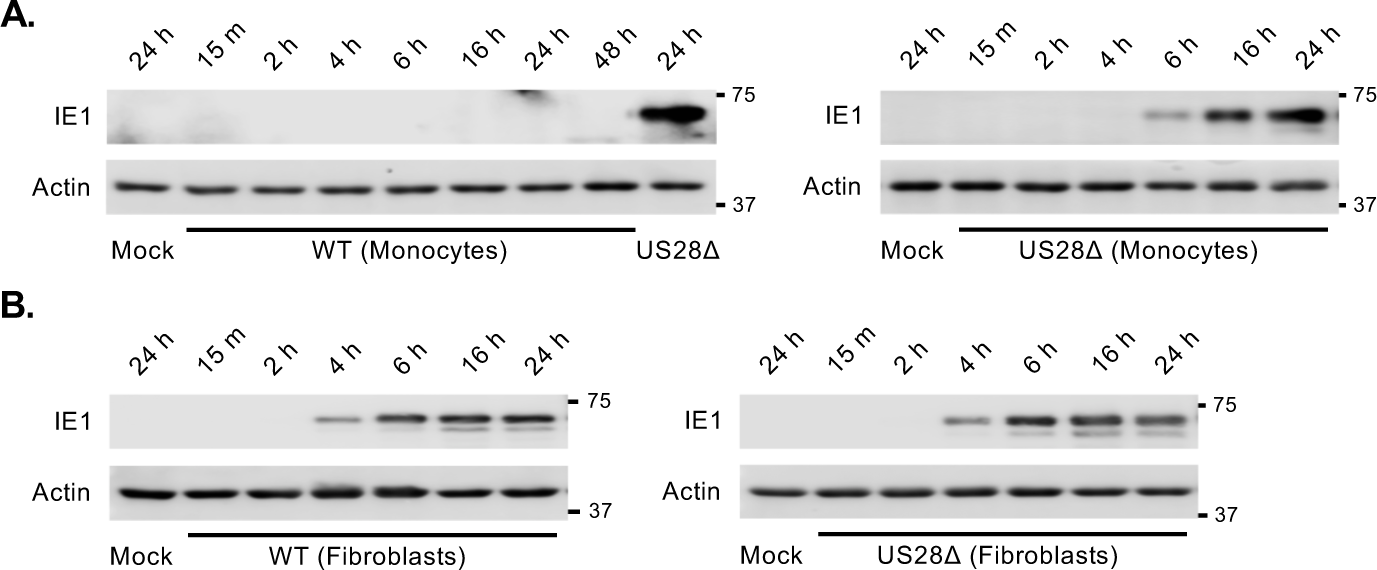
Kinetics of IE1 expression in US28Δ infected monocyte is similar to fibroblast infection. **(A)** Monocytes and **(B)** fibroblasts were infected (MOI = 1) with mock, WT, or US28Δ for the indicated times, and IE1 expression was detected by immunoblot. All membranes were probed for β-actin as a loading control. Western blots are representative of at least 3 independent experiments.

### Virion-associated US28 blocks early IE1 expression in monocytes

Virion-associated US28 is important for establishing latency in Kasumi-3 and CD34+ HPCs (*15*), while *de novo* synthesized US28 is required to maintain latency (*10, 12, 14–17*). Thus, we next assessed the kinetics of US28 expression following infection of primary monocytes. To this end, we infected primary monocytes with TB40/E*mCherry*-US28-3xF (US28-3xF), a viral recombinant that contains a triple FLAG epitope tag inserted in-frame with the US28 ORF virus at the C-terminus (*20*). We also included lytically-infected primary fibroblasts as a control. Although US28 expression was observed in fibroblasts at 48 hpi, we failed to detect *de novo* synthesized US28 in monocytes by western blot (**Fig. 3A**) or RT-qPCR (**Fig. 3B**) until 4 days post-infection (dpi) (**Fig. S4**). SV40-driven mCherry was detected in US28-3xF-infected monocytes by 24 hpi using western blot (**Fig. 3A**) and flow cytometry (**Fig. S5**), confirming delivery of the viral genome to the nucleus. These data indicate *de novo* synthesized US28 is unlikely to be involved in the rapid suppression of IE1 during infection of monocytes and that virion-delivered US28 is responsible for preventing IE1 expression. To test this possibility, monocytes were infected as described above and cell lysates collected as early as 15 minutes (min) post-infection over 24 h. Long exposure during image capture revealed delivery of virion-associated US28 to infected monocytes within 15 min of infection (**Fig. 3C**), which was reduced over 48 h through proteasomal degradation (**Fig. 3C; Fig. S6**). To determine if virion-associated US28 suppresses IE1 expression upon infection, we infected monocytes with US28-complemented US28Δ (US28comp) HCMV that lacks the *US28* gene but expresses virion-associated US28 protein derived from the complementing cell line. We found IE1 protein expression was suppressed in US28comp-infected monocytes (**Fig. 3D, 3E**), indicating virion-delivered US28 is sufficient to attenuate IE1 expression, which is consistent with previous findings (*15*). Altogether, our data suggest that virion-associated US28 suppresses IE1 expression in order to promote the establishment of HCMV quiescence within monocytes.

**Fig 3.**
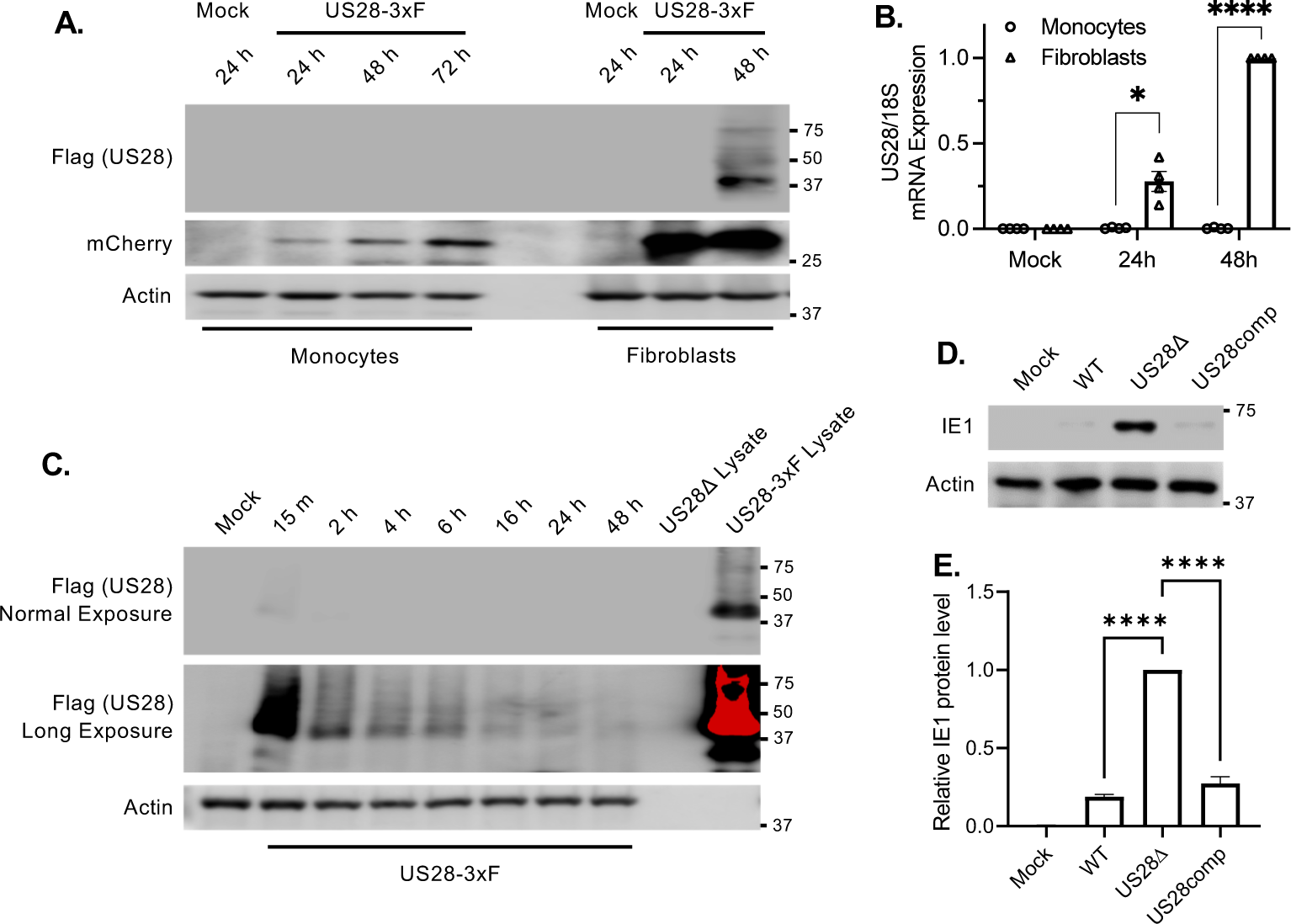
Virion-associated US28 inhibits IE1 expression. **(A, B)** Monocytes or fibroblasts were infected at MOI 1 or **(C)** MOI 5 (monocytes) for the indicated time with mock or US28-3xF. US28 protein and mRNA expression were detected by **(A, C)** western blot or **(B)** RT-qPCR, respectively. **(A, C)** SV40-driven mCherry expression in infected cells was assessed using an anti-mCherry antibody as a control for infection. US28-3xF and US28Δ cell-free virus lysates were used as positive and negative controls for US28, respectively. The red pixels represent the over saturation of the positive control band from long exposure of the blot during image capture. **(D, E)** Monocytes were infected (MOI = 1) with mock, WT, US28Δ, or US28-complemented US28Δ (US28comp) for 24 h and IE1 expression was detected by western blot. All membranes were probed for β-actin as a loading control. Western blots are representative of at least 3 independent experiments. Densitometry was performed using Image Lab software (Bio-Rad) from 3 independent blood donors. Statistical significance was measured using one-way ANOVA. ****P < 0.0001, *P<0.05.

### US28 modulates Akt activity to restrict *UL123* transcription

Activation of EGFR and its downstream pathways, such as phosphoinositide 3-kinase (PI3K) and Akt, are essential for the maintenance of HCMV latency, as inhibition of these proteins results in lytic replication of HCMV in CD34+ HPCs (*53, 54*). To investigate if activation of this pathway is regulated by US28 during the establishment of quiescent infection in monocytes and if inhibition leads to IE1 expression during WT infection, we infected monocytes with WT or US28Δ followed by treatment with small-molecule inhibitors targeting EGFR, PI3K, or Akt. However, we failed to detect IE1 expression in WT-infected monocytes in the presence of the inhibitors, suggesting that signaling from the EGFR/PI3K/Akt pathway induced during viral entry may play a distinct role in the early establishment of a quiescent infection in monocytes compared to the long-term maintenance of latency with CD34+ HPCs. In contrast to WT infection, US28Δ-infected monocytes treated with vehicle (DMSO) showed robust *UL123* transcription and IE1 protein expression, which was significantly attenuated in the presence of each inhibitor (**Fig. 4A to 4C**). Selected inhibitor concentrations did not affect cell viability (**Fig. S7A**) or viral entry (**Fig. S7B**), indicating signaling from the EGFR/PI3K/Akt pathway is necessary for early *UL123* transcription in US28Δ-infected monocytes. As the phosphorylation signature of Akt is responsible for controlling the activity levels (*40, 41*), and thus for its substrate specificity (*55–58*), we next assessed the activation profile of Akt in WT-versus US28Δ-infected monocytes. Consistent with our previous reports (*42, 43, 59*), WT infection of monocytes preferentially induced a site-specific phosphorylation of Akt at S473 during HCMV entry into monocytes, whereas T308 activity is unchanged relative to mock-infected control cells (**Fig. 4D, 4E**). Surprisingly, infection with US28Δ stimulated the phosphorylation of Akt at both S473 and T308 residues, which are required for the full activation of Akt and observed following canonical Akt activation by growth factors (*40, 41*). Moreover, the site-specific phosphorylation of Akt at S473 in US28comp-infected monocytes was similar to WT-infected cells (**Fig. S8**), indicating the importance of virion-associated US28 in dampening Akt activation during viral entry. Accordingly, proline-rich Akt substrate of 40 kDa (PRAS40), a known Akt substrate (*60*), was more robustly activated in US28Δ-infected cells when compared to WT-infected cells (**Fig. S9**). Similarly, mTORC1 activity, another downstream target of Akt, was enhanced in monocytes infected with US28Δ relative to WT-HCMV as evident by S6K phosphorylation (**Fig. S9**), a known marker for mTORC1 activation (reviewed in (*61*)). In addition to Akt, pUL38 has been shown to antagonize the ability of the tumor suppressor protein complex to negatively regulate mTORC1 and is also expressed in US28Δ-infected monocytes (**Fig. S1A**) (*62*). However, our data suggest that Akt activity is the major early regulator of mTORC1 in US28Δ-infected monocytes as loss of Akt activity prevented S6K phosphorylation (**Fig. S9**) and IE1 expression (**Fig. 4A**). Overall, these data suggest virion-associated US28 redirects Akt signaling during early infection to prevent the initiation of UL123 transcription and the establishment of HCMV quiescence in monocytes.

**Fig 4.**
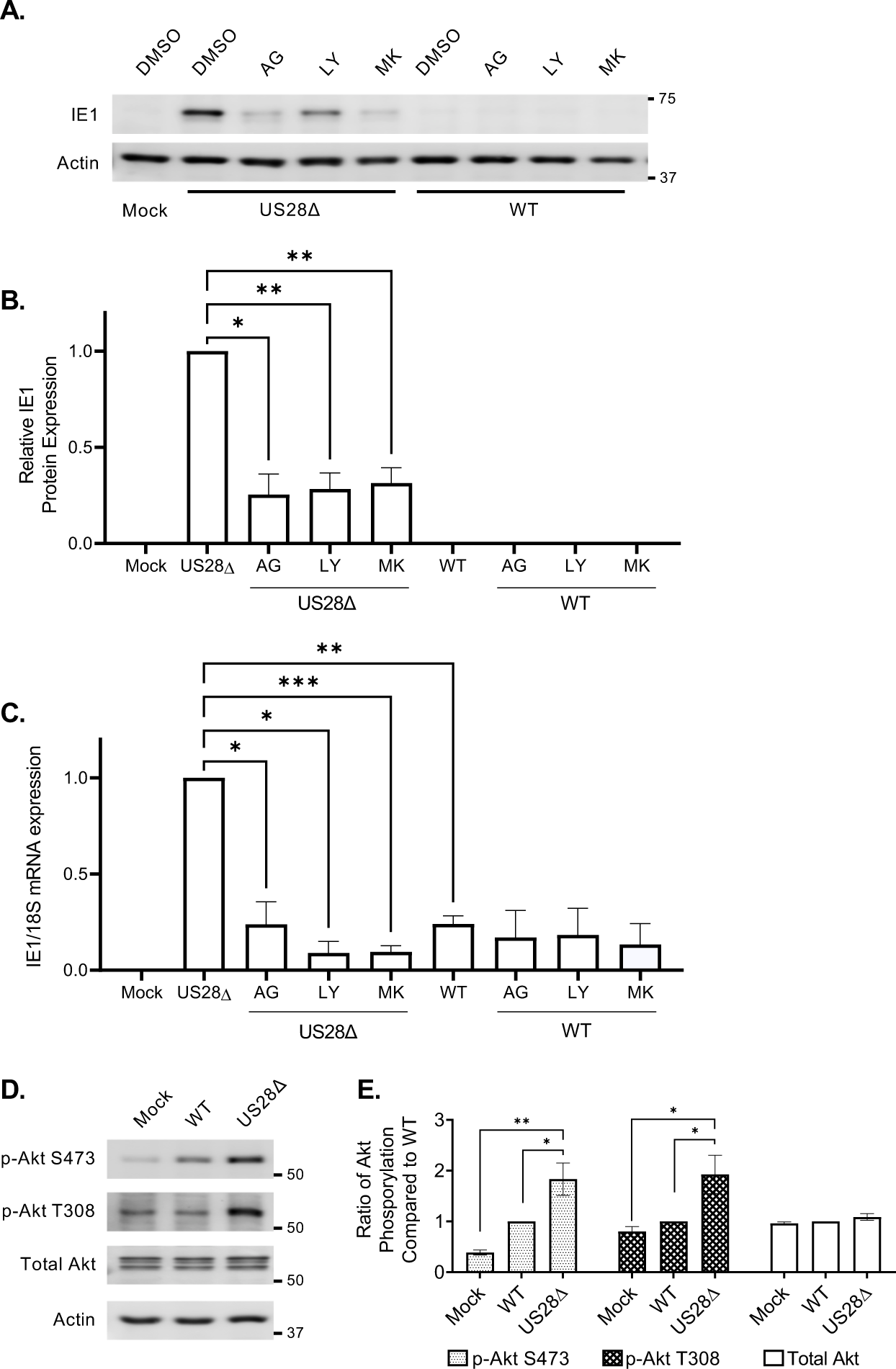
US28 modulates Akt pathway to establish quiescence. **(A to C)** Monocytes were infected with WT or US28Δ (MOI = 1). After 30 min (to allow uninterrupted viral entry), samples were treated with AG (5 µM; EGFR inhibitor), LY (25 µM; pan-PI3K inhibitor) or MK (10 µM; Akt inhibitor). After 24 h, cells were lysed, and IE1 expression was measured by **(A, B)** western blot and **(C)** RT-qPCR. **(D, E)** Monocytes were lysed 30 min post-infection, and specific antibodies were used to detect Akt activation. All membranes were probed for β-actin as a loading control. Western blots are representative of at least 3 independent experiments. Densitometry was performed using Image Lab software (Bio-Rad) from at least 3 independent blood donors. Statistical significance was measured using one-way ANOVA. ***P < 0.0005, **P < 0.005, *P<0.05.

### US28 dampens Akt activity induced by HCMV entry into monocytes to prevent IE1 expression

During canonical activation of Akt, receptor tyrosine kinases (RTKs) activate PI3K, which phosphorylates PI(4,5)P_2_ to PI(3,4,5)P_3_. PI(3,4,5)P_3_ then recruits Akt to the plasma membrane leading to the phosphorylation of Akt at T308 and S473 residues by PDK1 and mTORC2, respectively (*40, 41*). To examine if Akt phosphorylation at S473 and T308 regulates IE1 expression, we used several strategies to force the phosphorylation at both S473 and T308 during WT infection. First, we pretreated monocytes with GM-CSF prior to infection to stimulate Akt phosphorylation at both S473 and T308 (*63*). Indeed, we found GM-CSF-induced phosphorylation at S473 and T308 (**Fig. 5A**) corresponded with IE1 expression during WT infection (**Fig. 5B, 5C**). We did not observe an increase in macrophage-associated markers on the surface of WT-infected monocytes treated with GM-CSF by flow cytometry when compared to untreated WT-infected cells at 24 hpi (**Fig. S10**), suggesting the short duration of GM-CSF treatment is unlikely to promote sufficient monocyte differentiation that could result in IE1 expression in WT-infected cells. As a second approach, we added exogenous PI(3,4,5)P_3_ lipid (PIP3) to induce Akt S473 and T308 phosphorylation (**Fig. 5D**) (*40, 41*), which corresponded to increased IE1 expression in WT-infected monocytes (**Fig. 5E, 5F**). Surprisingly, the SH2 domain-containing inositol 5-phosphatase (SHIP) 1 product PI(3,4)P_2_ (PIP2), which is responsible for the site-specific phosphorylation of AKT at S473 during WT infection (*42, 43*), also led to IE1 expression, albeit to a lesser extent than PI(3,4,5)P_3_ treatment. However, IE1 expression induced by PI(3,4)P_2_ was likely due to the unexpected increase in T308 phosphorylation along with the expected increase in S473 phosphorylation (**Fig. 5D**). Finally, we transfected monocytes with an Akt-expressing construct where the N-terminus of Akt is fused to a myristylation domain (myr-Akt), which results in the recruitment of Akt to the cell membrane independent of its pleckstrin homology (PH) domain (*63, 64*). As a result, Akt is constitutively phosphorylated at both S473 and T308 residues by mTORC2 and PDK1, respectively (*56, 58*). As expected, WT-infected monocytes expressing the myr-Akt construct had robust phosphorylation at both S473/T308 residues (**Fig. 5G**). Of note, myr-Akt lacks the PH domain (amino acids 4-129) resulting in a lower molecular weight band than endogenous Akt. Nonetheless, myr-Akt phosphorylation at S473/T308 resulted in robust IE1 protein expression in WT-infected monocytes (**Fig. 5H, 5I**). Additionally, we found that transcription of *UL123* was significantly increased in WT-infected cells with myr-Akt (**Fig. S11**), indicating IE1 expression in WT-infected monocytes with Akt phosphorylation at S473/T308 is not due to enhanced Akt/mTORC1-mediated protein translation. Overall, our data argues that phosphorylation of Akt at both S473 and T308 sites during HCMV entry leads to the initiation of UL123 transcription and subsequent IE1 expression.

**Fig 5.**
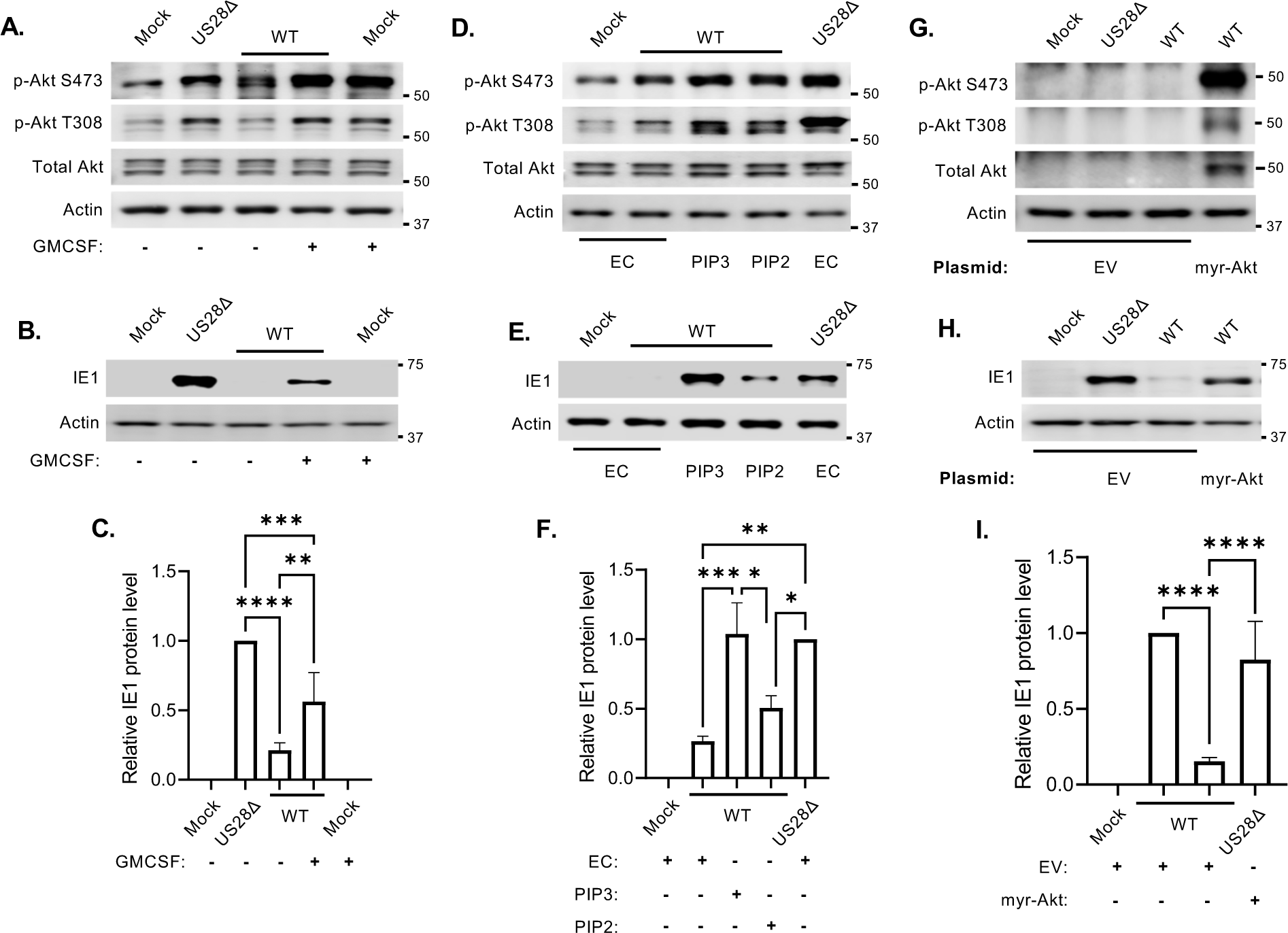
Induction of Akt T308 phosphorylation results in IE1 expression during WT infection. **(A to C)** Monocytes were treated with PBS or GM-CSF (100 ng/ml) for 30 min. Cells were then infected with mock, WT, or US28Δ (MOI = 1) for **(A)** 30 min or **(B, C)** 24 h**. (D to F)** Monocytes were treated with 15 µM of indicated phosphatidylinositol lipid (PIP) or empty lipid carrier (EC) for 50 min. Cells were next infected with WT and US28Δ for **(D)** 30 min or **(E, F)** 24 h. **(G to I)** Monocytes were transfected with myristylated-Akt (myr-Akt) plasmid or empty vector (EV). After 48 h, cells were mock, WT, or US28Δ infected for **(G)** 30 min or **(H, I)** 24 h. Expression of IE1, total Akt, phosphorylated (p) Akt S473 and p-Akt T308 were detected by western blot. All membranes were probed for β-actin as a loading control. Western blots are representative of at least 3 independent experiments. Densitometry was performed using Image Lab software (Bio-Rad) from 4 independent blood donors. Statistical significance was measured using one-way ANOVA. ****, P < 0.0001, ***P < 0.0005, **P < 0.005, *P<0.05. E= Endogenous Akt, M= Myristylated Akt, PIP3= PI(3,4,5)P_3_, PIP2= PI(3,4)P_2_.

Akt activity levels and substrate specificity depends on the phosphorylation ratio between S473 and T308 (*55–58*); thus, we asked whether the US28-mediated dampening of Akt activity (via the loss of T308 phosphorylation) induced during viral entry is necessary to prevent the initiation of IE1 expression. To this end, we knocked down PDK1 with siRNAs (**Fig. 6A**), which reduced Akt phosphorylation at T308, in US28Δ-infected monocytes (**Fig. 6B**). PDK1-deficient monocytes infected with US28Δ had reduced levels of IE1 compared to US28Δ-infected cells transfected with non-targeting control siRNA (**Fig. 6C, 6D**), indicating the reduction of T308 phosphorylation mediated by US28 is necessary to ensure that IE1 expression is not initiated during WT infection. Additionally, we transfected monocytes with siRNA targeting rictor (critical for the kinase activity of mTORC2) to assess if the full activation of Akt through phosphorylation of S473 together with T308 is required of IE1 expression (**Fig. 6A**). Rictor-deficient monocytes infected with US28Δ exhibited lower levels of S473 relative to cells transfected with a control siRNA (**Fig. 6B**). Importantly IE1 expression was significantly attenuated in rictor-deficient, US28Δ-infected monocytes (**Fig. 6C, 6D**), indicating that the full activation of Akt, via S473 and T308 phosphorylation, is necessary for IE1 expression. Collectively, our data indicates US28 dampens HCMV-induced Akt activity by preventing T308 phosphorylation. This partial Akt activity allows for the expression select antiapoptotic proteins necessary for the survival of HCMV-infected monocytes without initiating IE gene expression associated with fully active Akt.

**Fig 6.**
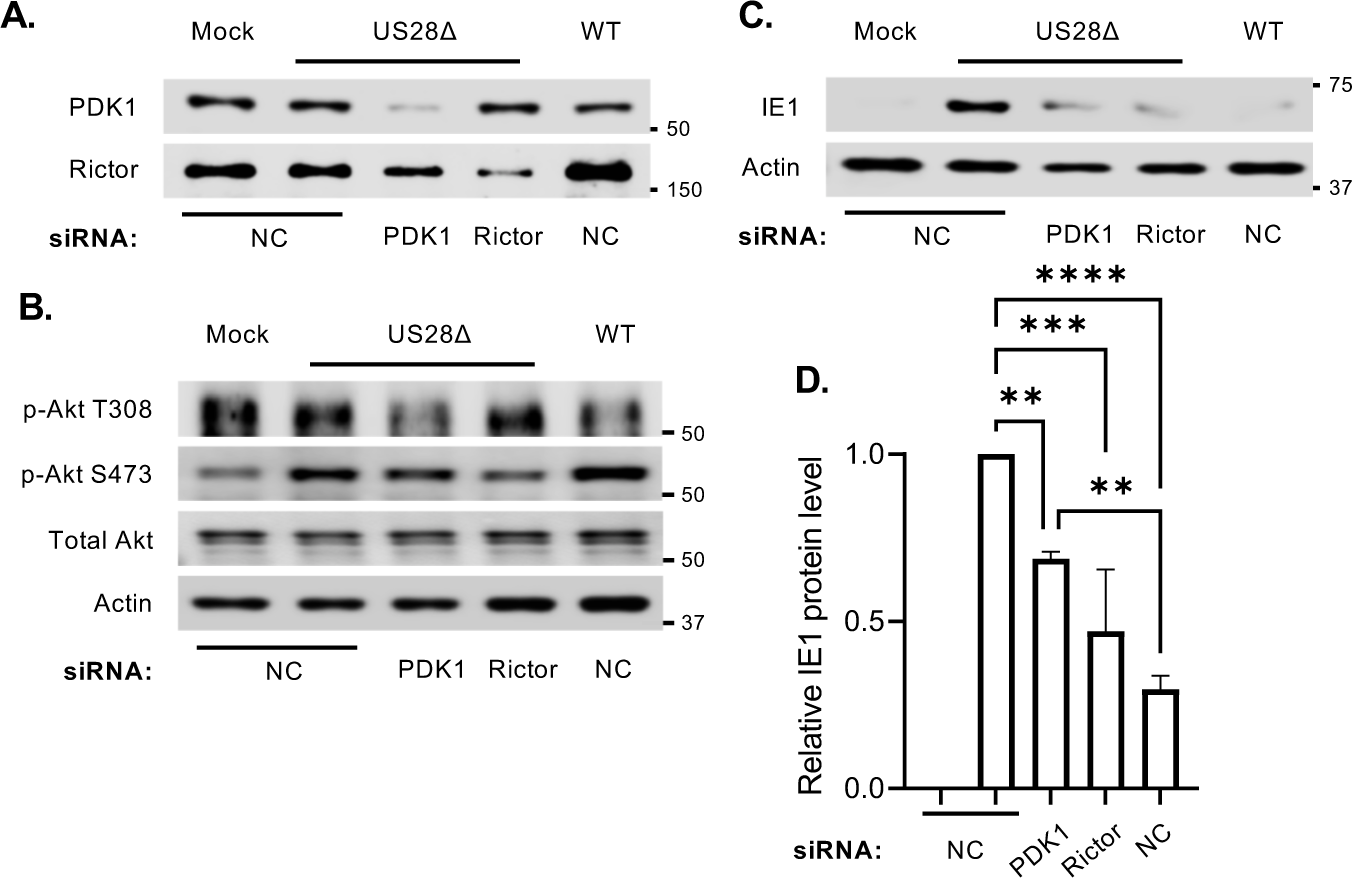
Akt phosphorylation at both S473 and T308 residues is required for IE1 expression in US28Δ-infected monocytes. Monocytes were transfected with scramble (NC), PDK1, or rictor siRNA and incubated for 48 h, after which cells were WT-or US28Δ-infected for **(A, B)** 30 min, or **(C, D)** 24 h. Levels of PDK1, rictor, total Akt, p-Akt S473, p-Akt T308, and IE1 were determined by western blot. All membranes were probed for β-actin as a loading control. Western blots are representative of at least 3 independent experiments. Densitometry was performed using Image Lab software (Bio-Rad) from 3 independent blood donors. Statistical significance was measured using one-way ANOVA. **** P < 0.0001, ***P < 0.0005, **P < 0.005.

### US28 restricts EGFR activation to regulate Akt phosphorylation

Next, we investigated the mechanism by which US28 diminishes Akt phosphorylation. During HCMV-mediated non-canonical activation of Akt, SHIP1 is activated and converts PI(3,4,5)P_3_ to PI(3,4)P_2_ leading to the site-specific phosphorylation of Akt at S473 (*42, 43*). Thus, we hypothesized that US28 promotes SHIP1 activity to redirect Akt phosphorylation towards S473 during WT infection. However, SHIP1 activity was similar between WT- and US28Δ-infected monocytes (**Fig. 7A, 7B**). Further, inhibition of SHIP1 using 3AC at concentrations that attenuated HCMV-induced S473 phosphorylation (**Fig. 7C**), but had no effect on viral entry or cell survival (**Fig. S12**), did not allow for *UL123* mRNA and IE1 protein expression during WT infection (**Fig. 7D to 7F**). These data argue that US28 may regulate factors upstream of SHIP1, such as EGFR. Indeed, we found that infection of monocytes with US28Δ enhanced EGFR phosphorylation when compared to WT infection (**Fig. 8A, 8B**). Additionally, inhibition of EGFR activity using AG1478 following US28Δ infection reduced Akt phosphorylation at both S473 and T308, compared to DMSO-treated infected cells (**Fig. 8C**), without affecting viral entry or monocyte viability at the concentrations used (**Fig. S7**). Similarly, inhibition of downstream PI3K resulted in decreased Akt phosphorylation (**Fig. 8C**), suggesting US28 modulates EGFR activation to finetune Akt activity. Moreover, infection with US28-recombinant mutants with defective signaling capabilities, ΔN and R129A, also resulted in increased EGFR activation and Akt phosphorylation at S473 and T308 (**Fig. 8D**). To assess if US28 is sufficient to limit EGFR activity, we utilize THP-1 cells, a monocytic cell line permissive for HCMV latency, transduced with a lentivirus construct (pSLIK-US28-3xF) that allows for the doxycycline (DOX) inducible expression of pUS28 fused to a C-terminal triple-FLAG epitope tag (*15*). The stable THP-1-pSLIK-hygro and THP-1-pSLIK-US28-3xF lines were generated as previously described (*15*). Indeed, we found expression of US28 alone attenuated EGF-induced EGFR activity (**Fig. 8E, 8F**). Together, these data indicate US28-mediated cell signaling restricts EGFR activation to limit HCMV-induced Akt activity during the establishment of HCMV quiescence in monocytes.

**Fig 7.**
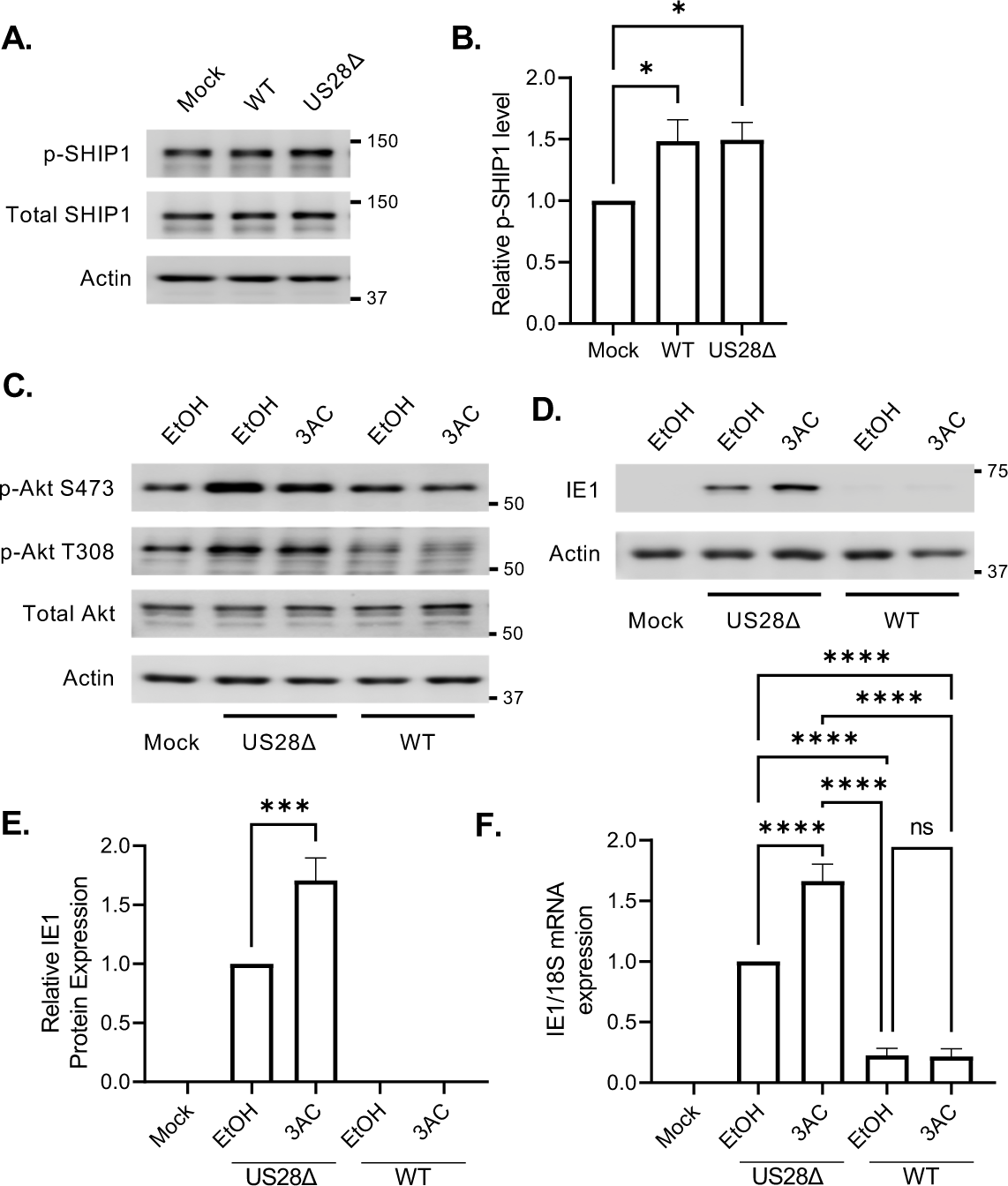
US28 does not regulate SHIP1 activity. **(A to F)** Monocytes were infected with mock, WT, or US28Δ (MOI = 1). After 30 min, cells were treated with ethanol (EtOH; carrier control) or 3AC (5 µM; SHIP1 inhibitor) and then incubated for **(A to C)** 30 min, or **(D to F)** 24 h. **(A to C)** Total SHIP1, p-SHIP1, total Akt, p-Akt S473, and p-Akt T308 were detected by western blot. IE1 expression was detected by **(D, E)** western blot or **(F)** RT-qPCR. All membranes were probed for β-actin as a loading control. Western blots are representative of at least 3 independent experiments. Densitometry was performed using Image Lab software (Bio-Rad) from 3 independent blood donors. Statistical significance was measured using one-way ANOVA.; ****, P < 0.0001, ***P < 0.0005, *P<0.05, ns= not significant.

**Fig 8.**
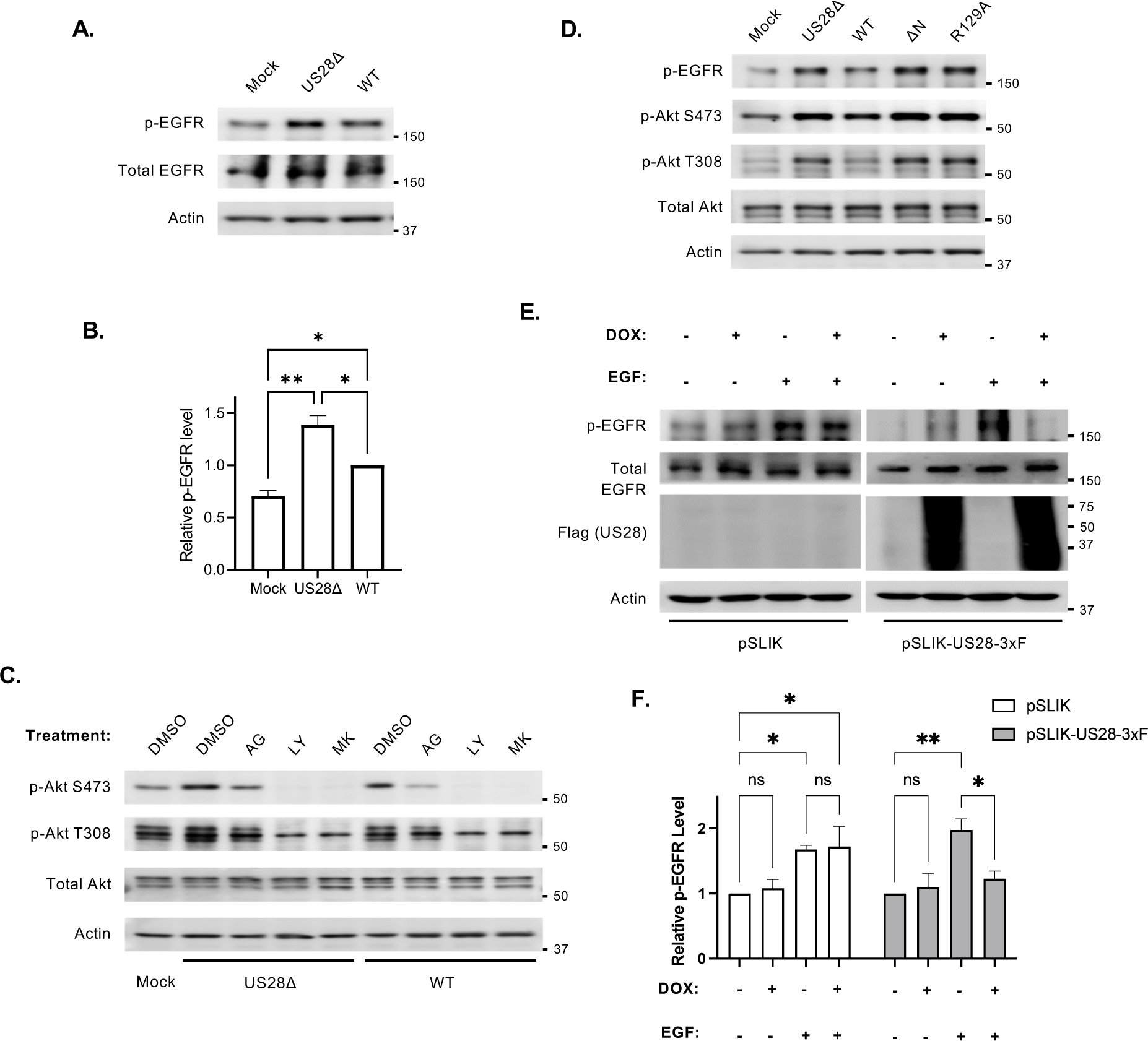
US28 limits HCMV-induced EGFR activation during viral entry. **(A to C)** Monocytes were mock, WT, or US28Δ infected (MOI = 1) for 30 min. **(A, B)** Cells were then immediately lysed and total EGFR and p-EGFR detected by western blot. **(C)** After 30 min of uninterrupted binding and entry, samples were treated with AG (5 µM), LY (25 µM), or MK (10 µM) for an addition 30 min. **(D)** Monocytes were infected (MOI = 1) with mock, WT, US28Δ, ΔN or R129A, for 30 min. **(C, D)** p-EGFR, total Akt, p-Akt S473, and p-Akt T308 were detected by western blot. **(E, F)** pSILK empty vector or pSLIK-US28-3xF-expressing THP-1 cells were treated with DOX for 24 hours. Cells were then treated with EGF (200 ng/mL) for 30 min. Levels of p-EGFR, total EGFR, and US28 (anti-flag antibody) were determined by western blot. All membranes were probed for β-actin as a loading control. Western blots are representative of at least 3 independent experiments. Densitometry was performed using Image Lab software (Bio-Rad) from 3 independent experiments. Statistical significance was measured using one-way ANOVA.; **P < 0.005, *P<0.05, ns= not significant.

## Discussion

Silent infections within the myeloid compartment play a critical role in HCMV’s dissemination (mediated by monocytes) and persistence (mediated by CD34+ cells) strategies. A significant effort has been made to elucidate the mechanisms regulating viral latency in myeloid cells (reviewed in (*65–67*)). However, mechanisms by which HCMV promotes the establishment of a quiescent infection within monocytes remain unclear. US28 is essential for latency/quiescence in CD34+ HPCs (*12, 14, 15, 17*), THP1 cells (*10, 15, 35*), Kasumi-3 cells (*12, 14, 15*), and monocytes (*10, 36, 37*); thus, we sought to determine the mechanism through which US28 promotes the establishment of a quiescent infection within monocytes. Herein, we demonstrate that virion-associated US28 is necessary and sufficient to dampen EGFR signaling to limit Akt activity triggered by HCMV entry. The reduced non-canonical activation is associated with a site-specific phosphorylation of Akt at S473 (**Fig. 9**), which we have shown upregulate a select subset of pro-survival proteins required for the long-term survival of normally short-lived monocytes (*42, 43, 59, 68*). In this study, we show the preferential phosphorylation at S473, but not T308, is also critical for the establishment of a quiescent infection, as the full activation of Akt by phosphorylation at both sites leads to the initiation of IE gene expression and subsequent lytic replication cycle **(Fig. 9)**. These results highlight virion-delivered US28 as a critical player to the viral dissemination strategy, as it rapidly modulates cellular signaling pathways triggered during viral entry to produce a cellular environment conducive to both the long-term survival of infected monocytes and the establishment of a quiescent infection.

**Fig 9.**
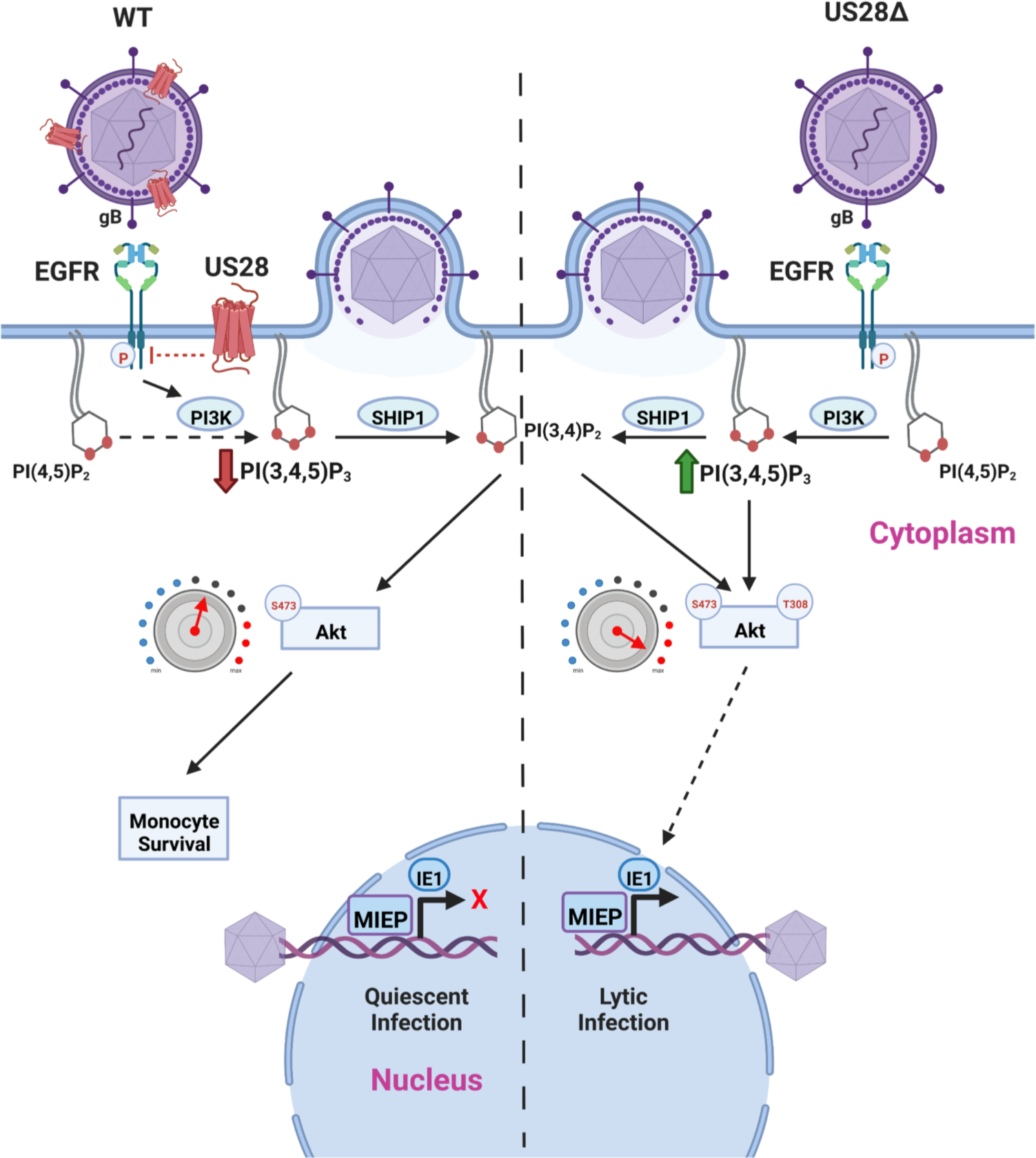
Proposed model for US28 regulation of quiescent HCMV infection. During WT HCMV infection of monocytes, US28 negatively regulates EGFR phosphorylation. Downstream of EGFR, PI3K phosphorylates PI(4,5)P_2_ to PI(3,4,5)P_3_. Additionally, HCMV infection results in increased SHIP1 activity, which dephosphorylates PI(3,4,5)P_3_ to PI(3,4)P_2_. Recruitment of Akt to PI(3,4)P_2_ leads to a preferential phosphorylation at S473 and establishment of a quiescent infection during WT infection. Contrarily, in US28Δ-infected monocytes, EGFR is robustly phosphorylated without impacting SHIP1 activity. Thus, in the absence of US28, an accumulation of PI(3,4,5)P_3_ leads to Akt phosphorylation at both S473 and T308, which is required for IE1 expression and lytic HCMV replication.

Akt is a central hub responsible for translating receptor-mediated signaling events into different cellular functional outcomes. HCMV has evolved to usurp Akt activity to support the different stages of the HCMV lifecycle. Initial studies found HCMV entry into permissive fibroblasts triggered an early activation of Akt, which was maintained throughout lytic replication to support viral entry, gene expression, and DNA replication (*69–72*). However, more recent studies indicate that AKT activity is suppressed as the lytic infection cycle progresses allowing for maximum virus production (*54, 73*). These findings underline a highly complex relationship between Akt and HCMV lytic infection, whereby Akt exerts both positive and negative effects dependent on the stage of the lytic replication cycle. During latent infection, PI3K/Akt signaling is also rapidly activated during the initial establishment of latency within CD34+ stem cells to promote viral entry (*74*). Following the establishment of latency, EGFR/PI3K/Akt signaling is required to maintain latency as inhibition of this pathway enhances viral reactivation from latently infected CD34+ cells (*54*). During a quiescent infection of monocytes, HCMV rapidly induces a noncanonical partial activation of Akt during entry of monocytes to promote the long-term survival of HCMV infected monocytes (**Fig. 4**) (*43, 75–77*). In contrast to latently infected CD34+ HPCs, inhibition of Akt did not lead to IE expression, but rather stimulating the full activation of Akt promoted the rapid expression of IE following viral entry into monocytes (**Fig. 5**). The discrepancy between the positive role of Akt on IE expression during the establishment of quiescence in monocytes versus the negative role of Akt during the maintenance of latency in CD34+ HPCs is unclear. The T308/S473 phosphorylation ratio of Akt was not examined in CD34+HPCs; thus, the possibility exists that Akt has a unique activation profile within monocytes. Alternatively, perhaps Akt activity changes as silent infections transition from establishment to maintenance. Regardless, how HCMV modulates the EGFR/PI3K/Akt signaling pathway and the subsequent functional outcomes appears to be highly cell specific and timing dependent.

Monocytes are short-lived cells programmed to undergo apoptosis 48 h after their release from the bone marrow in the absence of differentiation stimuli (*78–80*). Following stimulation with normal myeloid growth factors, Akt is activated by phosphorylation at both S473 and T308 residues to mediate the survival of short-lived monocytes (*81–83*). In contrast, HCMV promotes the survival of the infected monocytes by an atypical activation of Akt via the preferential phosphorylation at S473 (*42, 43*). The phosphorylation ratio between S473 and T308 dictates Akt activity as well as substrate specificity (*55–58*), suggesting that HCMV-activated Akt exhibits a distinct biological output compared to growth factor-activated Akt. Indeed, during quiescent infection of monocytes, HCMV upregulates a select subset of Akt-dependent, antiapoptotic proteins, including Mcl1, HSP27, and XIAP (*68, 84*). In this current study, we found that WT-mediated Akt phosphorylation at S473 did not result in IE expression, while US28Δ-mediated Akt phosphorylation at S473 and T308 led to IE1 production **(Fig. 5; Fig. 6B, 6C; Fig. 7C, 7D),** and the subsequent expression of viral early and late proteins **(Fig. S1)**. Because MIEP is differentially regulated by a multitude of transcription factors during lytic and latent infections (reviewed in (*66*)), it is attractive to speculate that S473 Akt does not activate the same subset of transcription factors as S473/T308 Akt. Our data would then suggest that US28 guides the precise activation of Akt to promote the survival of infected monocytes without triggering Akt-dependent transcription factors necessary for the initiation of IE gene expression. In support of this, US28 negatively regulates several transcription factors with known binding sites on the MIEP, including AP-1 and NFκB, which are required for attenuating IE gene expression and establishing latency within CD34+ (*15*), Kasumi-3 (*15*), and THP1 cells (*10*).

How US28 modulates the phosphorylation profile of Akt to generate a signaling network essential to the survival of infected monocytes, while also being beneficial to the establishment of quiescent infection, is unclear. During viral entry, the HCMV viral glycoprotein gB directly binds and activates EGFR (*42, 43*), which then activates downstream PI3K to phosphorylate PI(4,5)P_2_ lipid into PI(3,4,5)P_3_. During canonical EGFR/PI3K signaling, PI(3,4,5)P_3_ directs Akt phosphorylation at both S473 and T308. In naïve uninfected cells, PI(3,4,5)P_3_ can be dephosphorylated to PI(3,4)P_2_ by SHIP1 to negatively regulate Akt activity. However, in certain cancer cells, including leukemia cells, and HCMV-infected monocytes, the recruitment of Akt to PI(3,4)P_2_ leads to site-specific phosphorylation at S473 (*43, 85–87*). Thus, we hypothesized US28 enhances S473 phosphorylation by either increasing SHIP1 activity to promote the conversion of PI(3,4,5)P_3_ into PI(3,4)P_2_ or decreasing EGFR activity to reduce accumulation of PI(3,4,5)P_3_ within HCMV-infected monocytes. Our study demonstrates virion-associated US28 expression attenuates EGFR activity **(Fig. 8)** and has no impact on SHIP1 activity **(Fig. 7)**. As the conversion of PI(3,4,5)P_3_ to PI(3,4)P_2_ by SHIP1 appears to be the rate limiting step, we accordingly found the addition of exogenous PI(3,4,5)P_3_ during WT infection resulted in canonical S473 and T308 Akt phosphorylation **(Fig. 5D)**. These data suggest HCMV prevents the accumulation of PI(3,4,5)P_3_ by restricting EGFR activity, allowing PI(3,4)P_2_ to be the predominant signaling lipid responsible for mediating Akt phosphorylation.

Many biological effects of GPCRs occur through crosstalk with EGFR (*88, 89*). The mechanism by which virion-associated US28 regulates EGFR activity remains to be determined. One possibility is that US28 functions similar to cellular GPRC5A by binding and sequestrating EGFR, leading to its recycling/degradation through endocytic pathway (*90*). Another possibility is through the proteolytic cleavage and subsequent release of EGFR ligands by US28. GCPRs stimulate the release of EGFR ligands (*91–94*), which induce a conformational change in EGFR, resulting in dimerization and auto-phosphorylation of the tyrosine kinase domain (*95–97*). However, the stability of dimerization is dictated by ligand specificity, affecting the phosphorylation and signaling properties of EGFR (*98*). Perhaps US28 attenuates EGFR activity by desensitization through proteolytic cleavage of EGFR ligands such as epiregulin and epigen, which are expressed on the monocyte surface and promote a less robust EGFR activation relative to EGF (*98, 99*). Although elucidating the mechanism through which US28 modulates Akt activity remains an active avenue of research, our study clearly demonstrates US28-mediated suppression of EGFR signaling plays a pivotal role in shaping the Akt signaling network to promote the establishment of quiescent HCMV infection within monocytes.

To the best of our knowledge, this is the first study to demonstrate the effect of differential Akt phosphorylation status in the regulation of IE gene expression during the establishment of a quiescent infection within primary monocytes. Suppression of the initial lytic infection in monocytes is critical for long-term viral persistence within monocytes and HCMV’s dissemination. Altogether, our study demonstrates virion-associated US28 attenuates EGFR in order to dampen and redirect Akt activity into generating a cellular environment simultaneously conducive to both monocyte survival and quiescent infection. Unraveling the interplay between viral and cellular signaling factors could identify new antiviral targets aimed at preventing reactivation of HCMV from the myeloid compartment.

## Materials and Methods

### Human Peripheral Blood Monocyte Isolation

Peripheral blood was drawn from random donors by venipuncture, diluted in Roswell Park Memorial Institute medium (RPMI) 1640 (ATCC, Product #30-2001), and centrifuged through Histopaque 1077 (MilliporeSigma) to remove red blood cells and neutrophils. Mononuclear cells were collected and washed with saline to remove the platelets and then separated by centrifugation through a Percoll (GE Healthcare) gradient (40.48% and 47.70%). More than 90% of isolated peripheral blood mononuclear cells were monocytes, as determined by CD14-positive staining (*100*). Cells were washed with saline, resuspended in RPMI 1640 (ATCC, Product # 30-2001), and counted. All experiments were performed in the absence of human serum (unless mentioned otherwise) at 37°C in a 5% CO_2_ incubator. SUNY Upstate Medical University’s Institutional Review Board and Health Insurance Portability and Accountability Act guidelines for the use of human subjects were followed for all experimental protocols in our study (IRB#: 262458-19). Isolation of human peripheral blood monocytes was performed as previously described (*9, 75*).

### Virus Preparation and Infection

Bacterial artificial chromosome (BAC)-derived TB40/E*mCherry* (wild type; WT) (*52*), TB40/E*mCherry*-US28-3xF (US28-3xF) (*20*), TB40/E*mCherry*-US28Δ (US28Δ) (*20*), TB40/E*mCherry*-US28ΔN-3xF (ΔN) (*15*), TB40/E*mCherry*-US28-R129A-3xF (R129A) (*15*), and US28-complemented US28Δ (US28comp) (*15*) were previously described. All virus stocks were propagated on human embryonic lung (HEL) 299 fibroblasts (CCL-137, ATCC) of low passage (P7-15) in Dulbecco’s Modified Eagle medium (DMEM) (Lonza) with 2.5 μg/ml plasmocin (Invivogen) and 10% fetal bovine serum (FBS) (MilliporeSigma). Virus was purified from the supernatant when 100% cytopathic effect (CPE) was observed by ultracentrifugation (115000 x *g*, 65 minutes (min), 22°C) through a 20% sorbitol cushion to remove cellular contaminants and resuspended in RPMI 1640 medium (ATCC, Product # 30-2001). A multiplicity of infection (MOI) of 1 genome copy/cell was used for each experiment unless otherwise stated. Mock infection was performed by adding an equivalent volume of RPMI 1640 medium to monocytes.

To determine the incorporation of US28 in US28-R129A, and US28 ΔN into mature virions, primary newborn human fibroblasts (NuFF-1; GlobalStem) were infected with TB40/E*mCherry*-US28-3xF (US28-3xF), TB40/E*mCherry*-US28-R129A-3xF (R129A), or TB40/E*mCherry*-US28ΔN-3xF (ΔN), respectively (MOI = 1.0 TCID_50_/cell). Once 100% cytopathic effect (CPE) was observed, the media was collected, precleared two times by low-speed centrifugation (3,000 x *g*), and then viral particles were purified through a 20% sorbitol cushion (51,610 x *g*, 90 min, room temperature). Viral particles were then concentrated 100X in 10 mM Tris-Cl, pH8.0, 400 mM NaCl, and 10 mM EDTA, and one-fifth of this was used for immunoblot analyses. As controls for expression of US28 and its mutants, whole cell lysates were also collected, lysed, denatured (at 42°C), separated by SDS-PAGE, and assessed by western blot. US28 expression was assessed by probing for the FLAG epitope tag.

### Virus-Binding and Entry Assay

To assess viral binding, monocytes were incubated on ice for 1 h and then infected with WT or US28Δ for 2 h on ice to allow for viral binding. Monocytes were then washed twice with ice-cold 1X PBS to remove any unbound viral particles. As a negative control, another set of infected monocytes were treated with proteinase K (1.0 mg/ml, 30 m on ice) after the 2 h incubation and then washed twice with ice-cold 1X PBS to remove bound virus. DNA was extracted with QIAamp DNA mini kit (Qiagen) according to the manufacturer’s recommendations.

To quantify viral entry, monocytes were pre-treated with AG (10 µM; EGFR inhibitor) for 1 h at 37°C to block efficient viral entry into monocytes or with vehicle control (DMSO). Cells were then incubated on ice for 1 h to allow for equal particle binding and then infected with WT or US28Δ for an additional h on ice. Infected cultures were then shifted to 37°C for 2 h to allow for viral entry into monocytes. Finally, monocytes were treated with proteinase K (1.0 mg/ml, 5 min at 37°C) and then washed twice with 1X PBS to remove the viral particles that remained outside of the cell membrane and were unable to enter monocytes. RNeasy mini kit (Qiagen) was used per the manufacturer’s recommendation to extract mRNA. Any contaminating DNA from the samples was then removed using TURBO DNA free kit (ThermoFisher), according to manufacturer’s protocols.

### Compound Treatments

Where indicated, the following reagents were used to treat the cells at concentrations noted in the text: Bafilomycin (Baf; inhibitor of lysosomal degradation of proteins (*101*)), MG132 (MG; proteasome inhibitor (*102*)), AG-1478 (AG; EGFR inhibitor (*103*)), 3-α-aminocholestane (3AC; SHIP1 inhibitor (*87*)) and LY294002 (pan-PI3K inhibitor (*104*)) all from Calbiochem; MK-2206 (MK; Akt inhibitor (*105*)) and staurosporine (ST; broad spectrum protein kinase inhibitor/inducer of apoptosis (*106*)) both from Selleckchem. For induction of canonical Akt activation, monocytes were treated with PBS or GM-CSF (100 ng/ml) for 30 min before infection. Additionally, where indicated, samples were treated with 15 µM of phosphatidylinositol lipids (PI(3,4,5)P_3_, Echelon Biosciences, Product #P-3908; PI(3,4)P_2_, Echelon Biosciences, Product #P-3416) or empty lipid carrier (EC; Echelon Biosciences, Product #P-9C1) for 50 min prior infection according to the manufacturer’s recommendations.

### Western Blot Analysis

Cells were harvested in modified radioimmunoprecipitation assay (RIPA) buffer (50 mM Tris-HCl [pH 7.5], 5 mM EDTA, 100 mM NaCl, 1% Triton X-100, 0.1% SDS, 10% glycerol), supplemented with protease inhibitor cocktail (MilliporeSigma) and phosphatase inhibitor cocktails 2 and 3 (MilliporeSigma) for 30 min on ice. The lysates were cleared from the cell debris by centrifugation at 4°C (5 min, 21000 x *g*) and stored at −20°C until further analysis. Protein samples were solubilized in Laemmli SDS-sample non-reducing (6x) buffer (Boston Bioproducts), supplemented with β-mercaptoethanol (Amresco) by incubation at 95°C for 5 min, unless otherwise stated. For detecting EGFR, phospho-EGFR and US28, protein samples were incubated at room temperature for 30 min after solubilizing in Laemmli SDS-sample nonreducing (6x) buffer (*107*). Equal amounts of total protein were loaded, separated by SDS-polyacrylamide gel electrophoresis (SDS-PAGE), and transferred to polyvinylidene difluoride membranes (Bio-Rad). Blots were blocked in 5% bovine serum albumin (BSA; Fisher Scientific) for 1 hour (h) at room temperature (RT) and then incubated with primary antibodies overnight at 4°C. The blots were then incubated with horseradish peroxidase (HRP)-conjugated secondary antibodies for 30 min at room temperature (RT), and chemiluminescence was detected using the Clarity Western ECL substrate (Bio-Rad). For details on antibodies used in this study, see Table S1. Densitometry was performed using Image Lab software (Bio-Rad).

### Semiquantitative and Quantitative PCR

Total DNA and mRNA were isolated using QIAamp DNA mini kit and RNeasy mini kit (Qiagen, Germantown, MD), per the manufacturer’s recommendation. TURBO DNA free kit (ThermoFisher, Waltham, MA) was used to remove the contaminating DNA from the extracted mRNA samples. Semiquantitative PCR was used to detect the expression of *UL122* (sense, 5’-CGCCTTCGTTACAAGCATCG-3’; antisense, 5’-AAGAGCAAACGCATCTCCGA-3’), *GAPDH* (sense, 5’-ACCCACTCCTCCACCTTTGAC-3’; antisense, 5’-CTGTTGCTGTAGCCAAATTCGT-3’) using Green GoTaq master mix (Promega, Madison, WI) with C1000 Touch Thermal Cycler (Bio-Rad, Hercules, CA). For quantitative PCR, *UL123* (sense, 5’-AGTGACCGAGGATTGCAACG-3’; antisense, 5’-CCTTGATTCTATGCCGCACC-3’), *US28* (sense, 5’-CCAGAATCGTTGCGGTGTCTCAGT-3’; antisense, 5’-CGTGTCCACAAACAGCGTCAGGT-3’) and *GAPDH* were detected with CFX Connect Real Time System (Bio-Rad) using iTaq Universal SYBR Green Supermix (Bio-Rad). For quantitative real time PCR (RT-PCR), iTaq Universal SYBR Green One-Step Kit was used to detect the expression of *UL123*, *US28* and 18S rRNA (sense, 5’-GCAATTATTCCCCATGAACG-3’; antisense, 5’-GGGACTTAATCAACGCAAGC-3’). For quantitative PCR, samples were analyzed in technical duplicate and normalized to GAPDH (for DNA) or 18S rRNA (for mRNA).

### Flow Cytometry

Monocytes were washed in 1X phosphate-buffered saline (PBS) and incubated in blocking solution, consisting of fluorescence-activated cells sorting (FACs) buffer (1X PBS, 2mM EDTA, 0.5% BSA), 5% bovine serum albumin (BSA), and human Fc receptor (FcR) binding inhibitor (eBioscience). Cells were stained with an allophycocyanin (APC)-anti-CD14 or APC-anti-mouse IgG1 isotype control antibody (BioLegend) on ice, then washed and stained with FITC-annexin V and propidium iodide (PI; ThermoFisher Scientific) to detect dead and dying cells. For differentiation studies, cells were stained with anti-CD86 and anti-CD163. For details on antibodies used in this study, see Table S1. After staining, cells were analyzed by flow cytometry using an LSRFortessa cell analyzer and BD FACSDiva software (BD Biosciences). For measuring mCherry expression, monocytes were washed in 1X PBS and resuspended in FACs buffer prior to analysis by flow cytometry.

### Plasmid and siRNA Transfection

Primary monocytes (3 × 10^6^ cells/transfection) were washed with 1X PBS and resuspended in 100 μl of P3 Primary Cell Nucleofector solution (Lonza) containing either 1000 ng empty vector (EV) (Addgene plasmid # 10841 (*108*)), or myristylated-Akt (myr-Akt) plasmid (Addgene plasmid # 26453 (*64*)). For knockdown of PDK1 and Rictor, 1000 nM validated Silencer Select siRNAs against PDK1 (Invitrogen-Themo Fisher Scientific; siRNA ID # s10274), Rictor (Invitrogen-Themo Fisher Scientific; siRNA ID # s48410), or Silencer negative-control (NC) siRNAs (Themo Fisher Scientific; catalog # AM4642) were mixed in the nucleofector solution prior to transfection. Plasmids and siRNAs were then transfected with a 4D-Nucleofector (Lonza) using program EI-100. Following transfection, monocytes were incubated in RPMI 1640 supplemented with 2% human AB serum at 37°C for 48 hours (h), after which transfected monocytes were infected (multiplicity of infection MOI = 1) with mock, WT, or US28Δ for 24 h. Whole cell lysates were collected and subjected to western blot analyses.

### Statistical analyses

All experiments were performed with a minimum of 3 biological replicates using primary monocytes isolated from different blood donors. Data were analyzed using ANOVA and Student’s t test comparison with GraphPad Prism software, and *p*-values less than 0.05 were considered statistically significant.

## Acknowledgments

We thank Christine Burrer in the Department of Microbiology and Immunology at SUNY Upstate Medical University for technical support, maintenance of lab operations, and assistance with virus growth and isolation. Portions of the paper were developed from the thesis of J. Mahmud. This work was supported by grants from the Carol M. Baldwin Breast Cancer Research Fund to G.C. Chan, National Institute of Allergy and Infectious Disease (R01AI141460 to G.C. Chan; R01AI153348 to C.M. O’Connor), and National Heart, Lung, and Blood Institute (R01HL139824) to G.C. Chan.

## Supplementary Information

**Fig S1.**
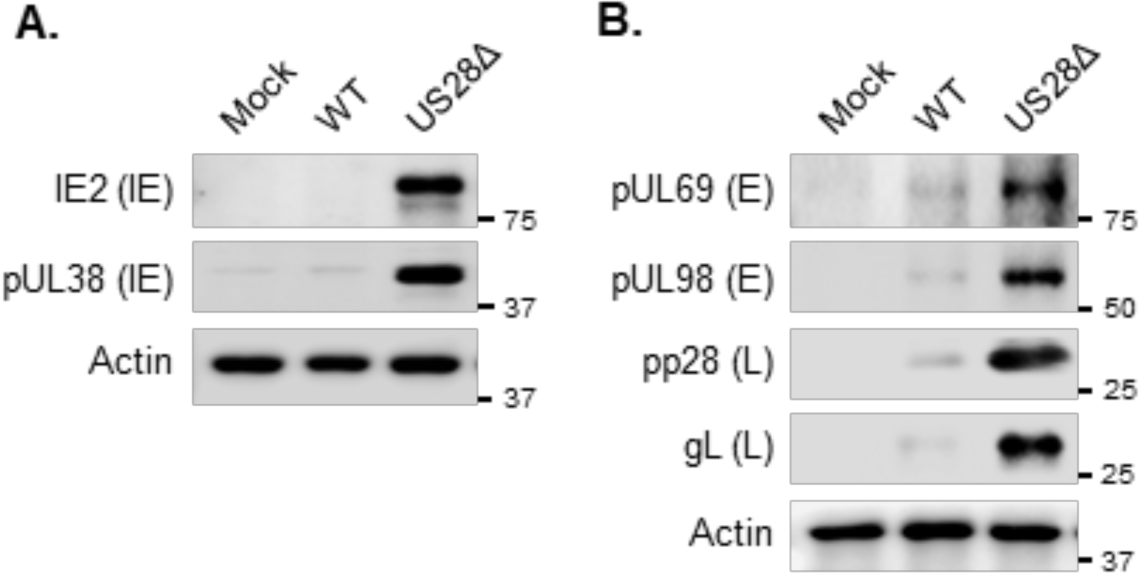
US28-mediated signaling prevents IE, E and L protein expression in monocytes. Monocytes were infected with mock, wild-type (WT), or US28Δ for **(A)** 24 h (MOI = 1), or for **(B)** 48 h (MOI = 5). Expression of immediate-early (IE), early (E) and late (L) proteins were detected by western blot. Membranes were probed for β-actin as a loading control. Representative blots are from 3 independent experiments.

**Fig S2.**
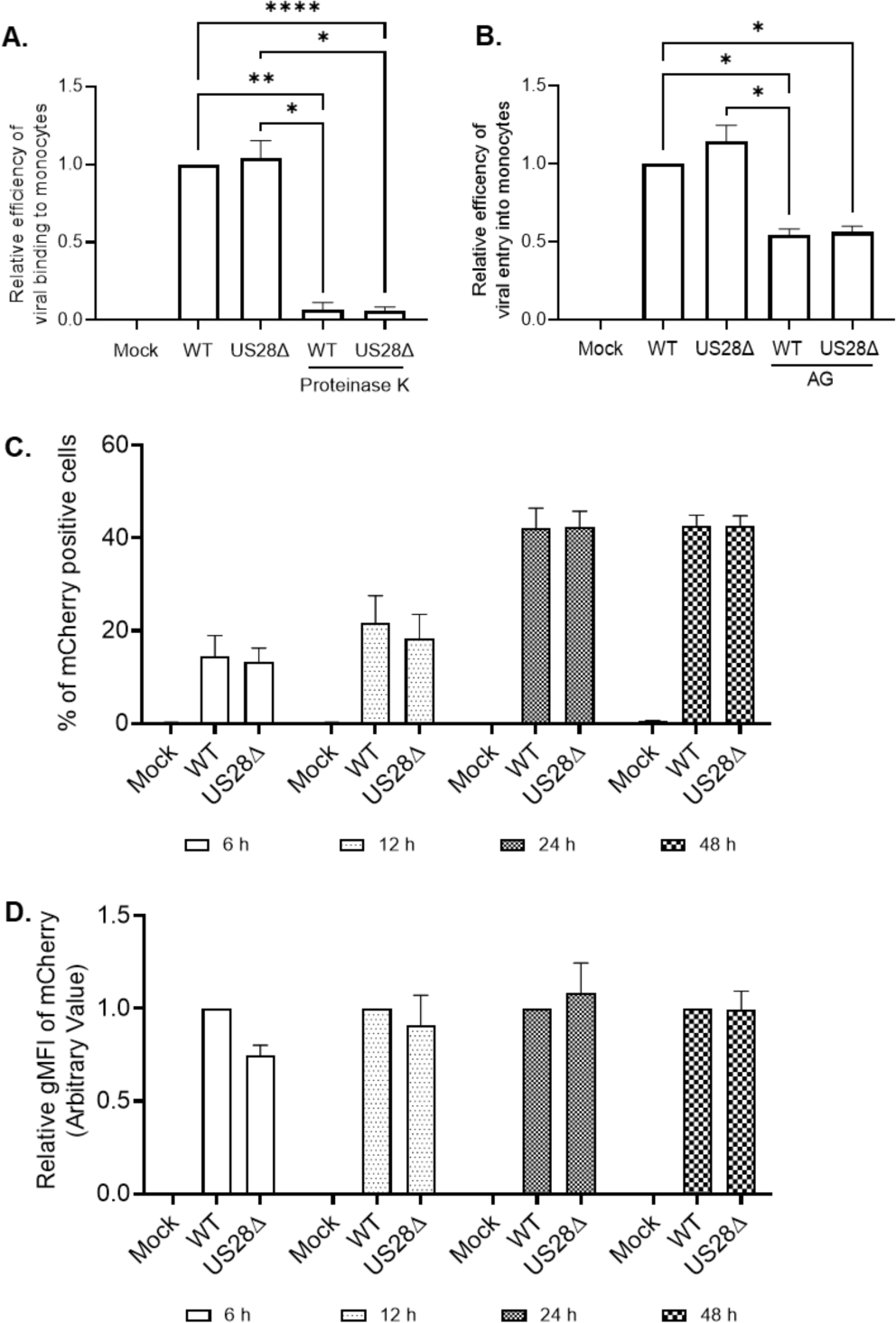
Deletion of the US28 ORF does not interfere with infection of monocytes or delivery of the viral genome to the nucleus. **(A)** Monocytes were incubated on ice for 1 h and then infected (MOI = 1) with WT or US28Δ for an additional 2 h on ice to allow for viral binding. Monocytes were then washed twice with ice-cold PBS to remove any unbound viral particles. As a negative control, another set of infected monocytes were treated with proteinase K (1.0 mg/ml, 30 min on ice) after the 2 h incubation and then washed twice with ice-cold PBS to remove the bound virus. **(B)** Monocytes were pre-treated with DMSO (vehicle control) or AG (10 µM; EGFR inhibitor, to block viral entry) for 1 h at 37°C. Cells were then incubated on ice for 1 h and then infected (MOI = 1) with WT or US28Δ for an additional 1 h on ice to allow viral binding. Infected monocytes were then shifted to 37°C for an additional 2 h to allow viral entry into monocytes. As a control, monocytes were treated with proteinase K (1.0 mg/ml, 5 min at 37°C) and then washed twice with PBS to remove viral particles that remained outside of the cell membrane and were unable to enter monocytes. **(A, B)** Samples were lysed and the viral genome was detected using RT-qPCR (UL123/GAPDH). **(C, D)** Monocytes were infected (MOI = 1) with mock, WT, or US28Δ for the indicated times. Expression of mCherry was detected using flow cytometry. Statistical significance was measured using one-way ANOVA; ****P < 0.0001, **P < 0.005, *P<0.05.

**Fig S3.**
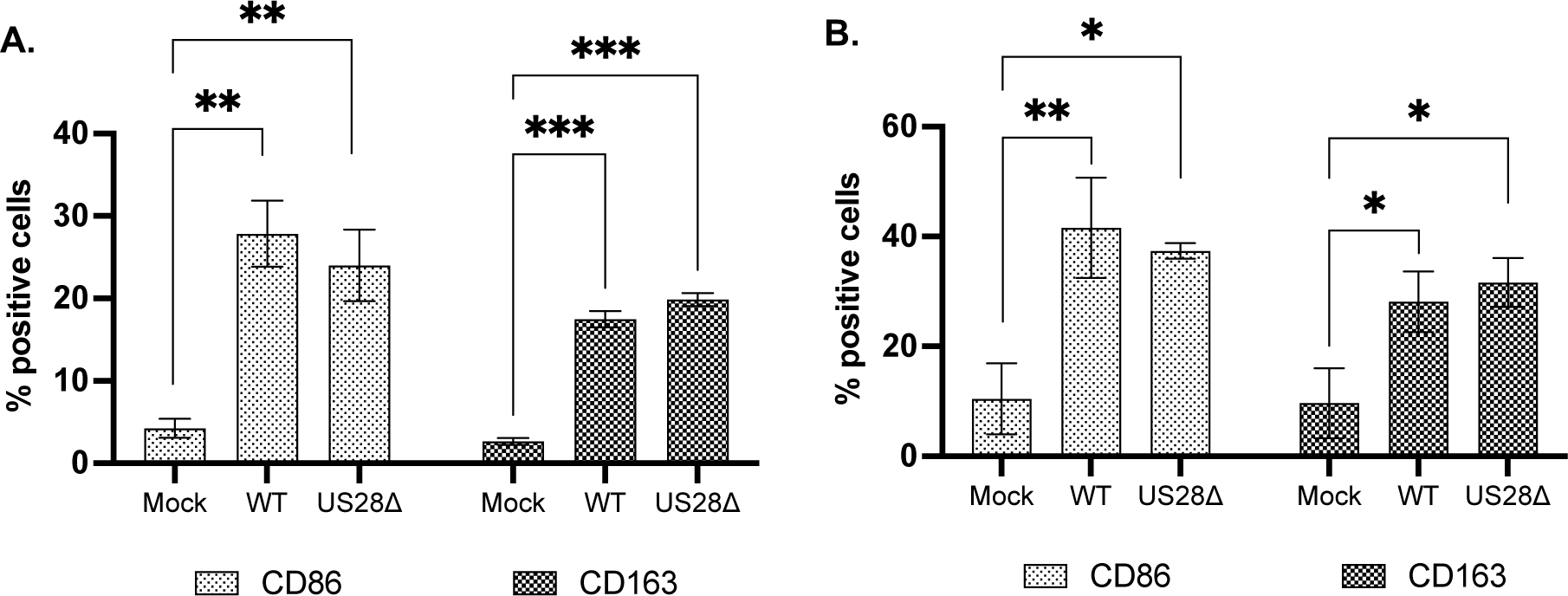
Deletion of the US28 ORF does not affect differentiation of HCMV-infected monocytes. Monocytes were infected (MOI = 5) with mock, WT, or US28Δ and incubated for **(A)** 24 h or **(B)** 7 days. The percent of positive cells for different macrophage markers were measured by flow cytometry. Statistical significance was measured using one-way ANOVA; ***P < 0.0005, **P < 0.005, *P<0.05.

**Fig S4.**
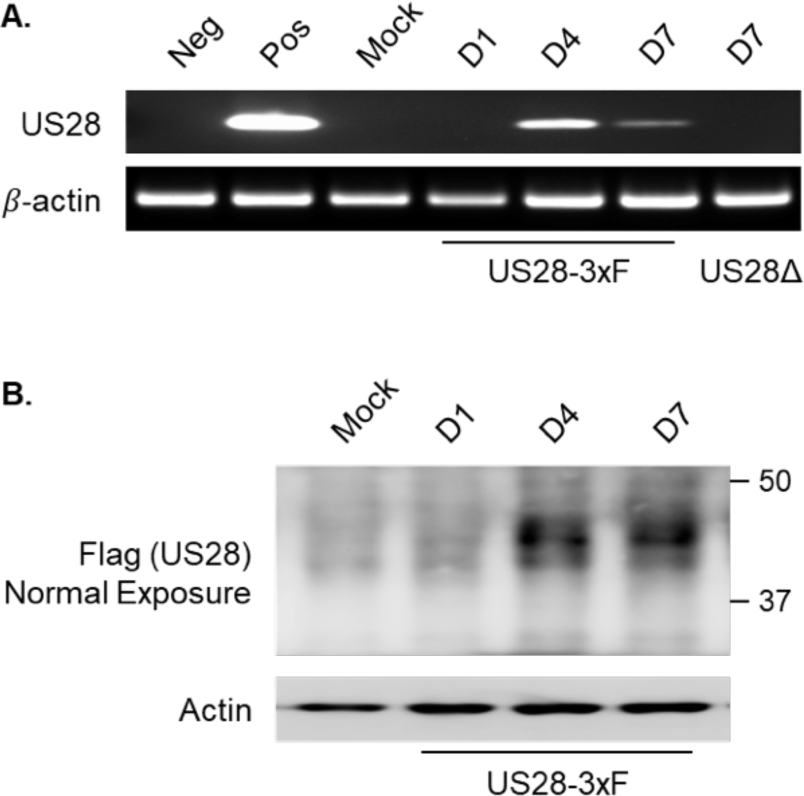
*De novo* synthesis of US28 is detected by 4 days post-infection. **(A, B)** Monocytes were infected (MOI = 1) with mock, US28Δ, or US28-3xF for 1, 4, or 7 days (D1, D4, D7). **(A)** Cells were lysed and expression of US28 and β-actin was detected by PCR. Uninfected and WT-infected fibroblasts (3 days post-infection) were used as negative (Neg) and positive controls (Pos), respectively. **(B)** Cells were lysed after the specified time and expression of US28 protein was detected using an anti-FLAG antibody by western blot. Membranes were probed for β-actin as a loading control. All results are representative of 3 independent experiments.

**Fig S5.**
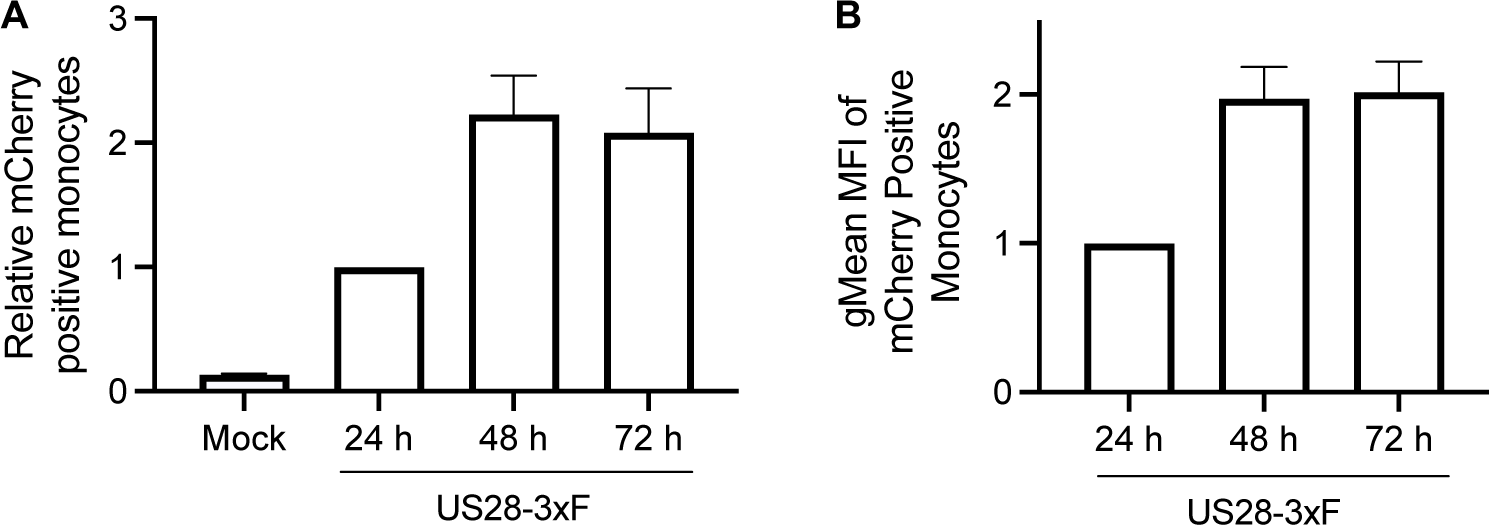
US28-3xF-infected monocytes express mCherry, which is detectable by flow cytometry. **(A, B)** Monocytes were infected (MOI = 1) with mock or US28-3xF for the indicated times. Expression of SV40-driven mCherry was then analyzed using flow cytometry.

**Fig S6.**
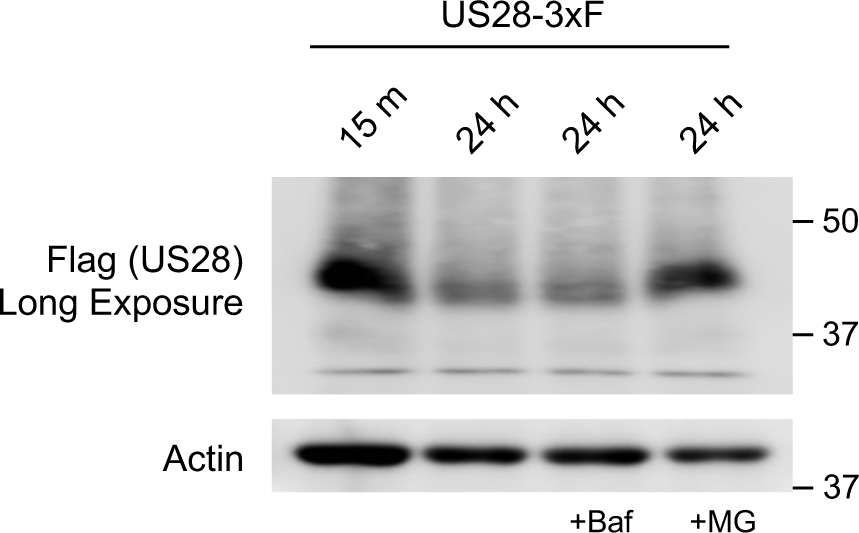
Virion-associated US28 is rapidly degraded by the proteasomal pathway in monocytes. Monocytes were pre-treated with Bafilomycin (200 nM) or MG132 (5 µM) for 1 h to inhibit lysosomal and proteasomal degradation, respectively. Cells were then infected (MOI = 1) with US28-3xF for the indicated times. Total cell lysates were analyzed for US28 expression by western blot. Membranes were probed for β-actin as a loading control. Results are representative of 3 independent experiments.

**Fig S7.**
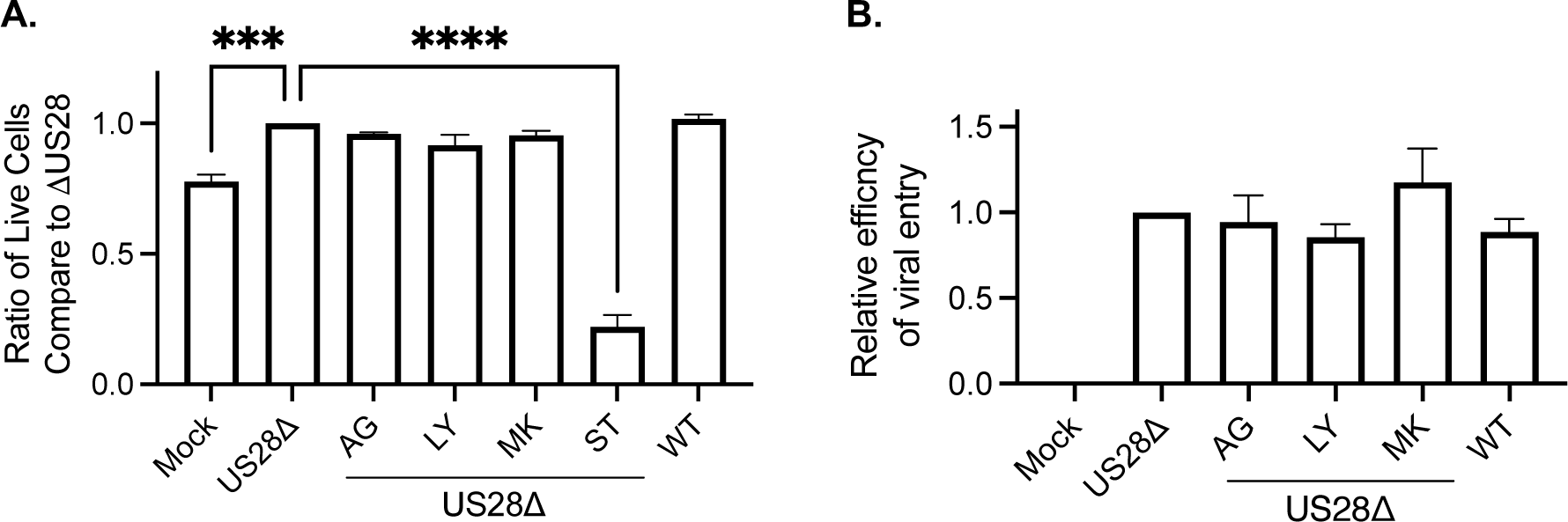
Selected concentrations of inhibitors of the EGFR/PI3K/Akt pathway does not affect survival of infected monocytes or viral entry. **(A, B)** Monocytes were infected (MOI = 1) with WT or US28Δ for 30 min to allow for viral entry. Where indicated, samples were then treated with AG (5 µM), LY (25 µM; pan-PI3K inhibitor), or MK (10 µM; Akt inhibitor). **(A)** After 24 h, viability of monocytes was measured by flow cytometry using annexin V and propidium iodide (PI) staining. Staurosporine (0.5 µM, ST; non-selective protein kinase inhibitor) was used as a positive control for cell death. **(B)** After 2 h, monocytes were treated with proteinase K (1.0 mg/ml, 5 min at 37°C) and then washed twice with PBS to remove the viral particles that remained outside of the cell membrane. Samples were then lysed, and the viral genome was detected using RT-qPCR (UL123/GAPDH). Statistical significance was measured using one-way ANOVA; ****P < 0.0001, ***P < 0.0005.

**Fig S8.**
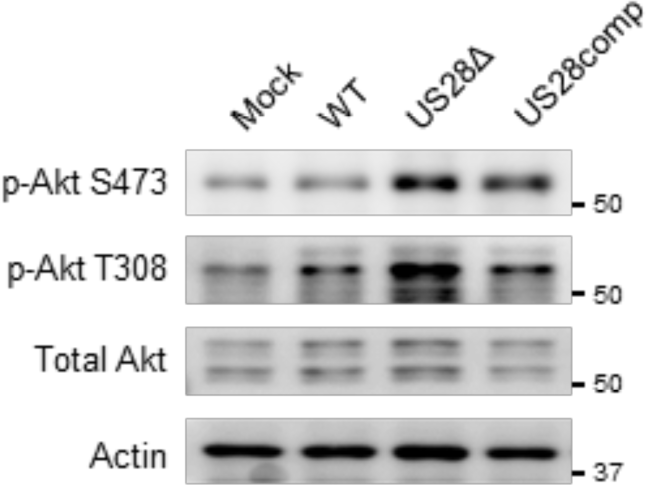
Virion-associated US28 suppresses Akt activation during HCMV entry into monocytes. Monocytes were infected (MOI = 1) with mock, WT, US28Δ, or US28comp for 30 min. Total Akt, p-Akt S473, and p-Akt T308 were detected by western blot. Membranes were probed for β-actin as a loading control. Results are representative at least 3 independent experiments.

**Fig S9.**
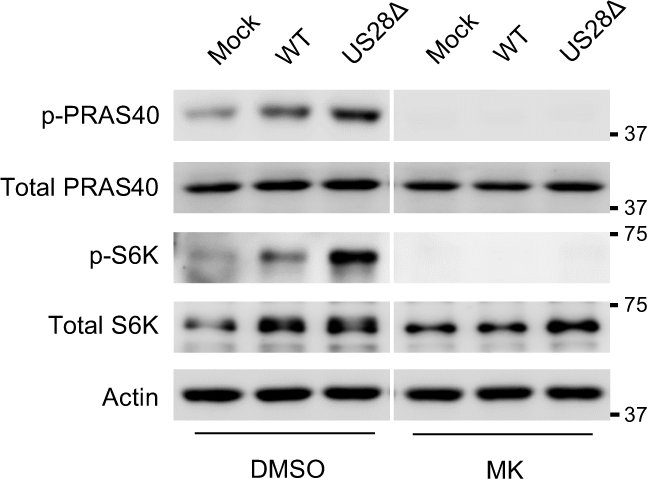
US28 dampens Akt activity following infection of monocytes. Monocytes were infected (MOI = 1) with mock, WT, or US28Δ. At 30 min post infection, samples were treated with DMSO or MK (10 µM). Monocytes were then incubated for an additional 30 min. Total PRAS40, p-PRAS40, p-S6K, and total S6K were detected by western blot. Membranes were probed for β-actin as a loading control. Results are representative of 3 independent experiments.

**Fig S10.**
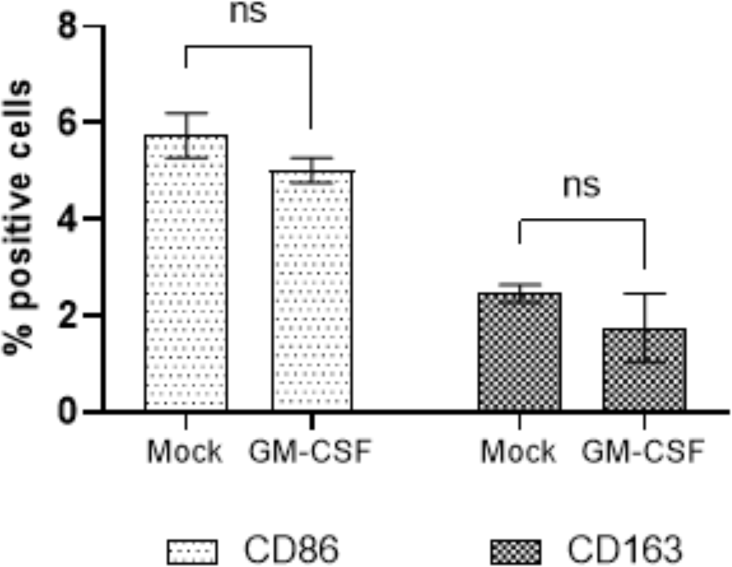
GM-CSF treatment in WT-infected monocytes does not enhance monocyte differentiation. Monocytes were treated with GM-CSF (100 ng/ml) and incubated for 24 h. The percent of positive cells for different macrophage markers were measured by flow cytometry. Statistical significance was measured using one-way ANOVA. ns= not significant.

**Fig S11.**
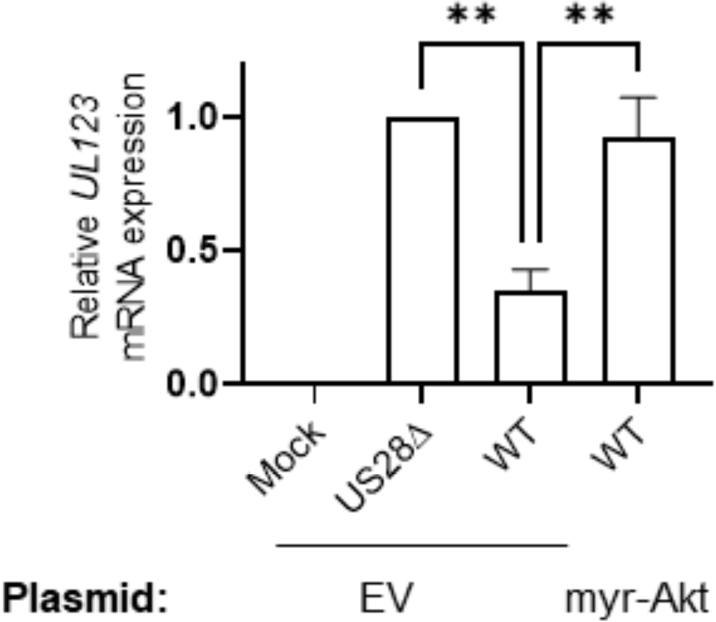
Expression of myr-Akt plasmid increases transcription of *UL123* in WT-infected monocytes. Monocytes were transfected with myr-Akt plasmid or EV for 48 h. Cells were then mock, WT, or US28Δ infected for 24 h and expression of IE1 mRNA was measured by RT-qPCR. Statistical significance was measured using one-way ANOVA. **P < 0.005.

**Fig S12.**
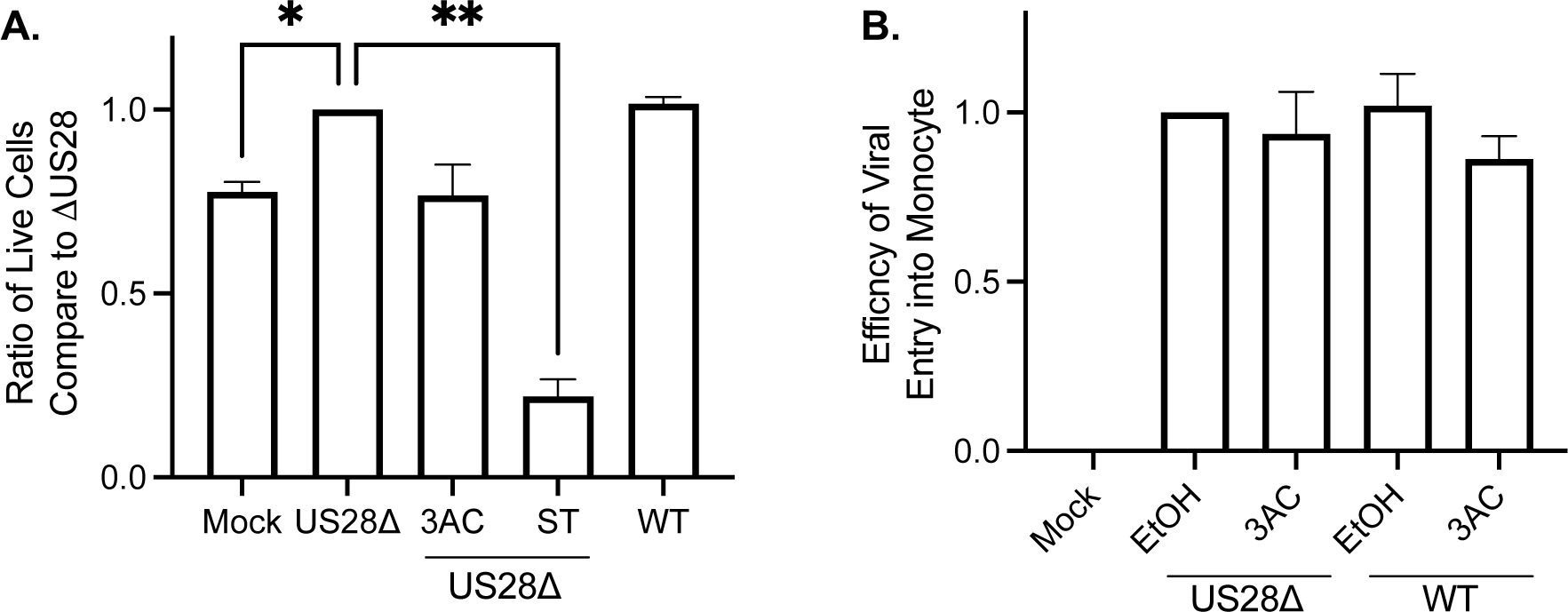
Selected concentration of SHIP1 inhibitor does not affect viral entry or survival of infected monocytes. **(A, B)** Monocytes were infected (MOI = 1) with WT or US28Δ for 30 min to allow for viral entry and then the indicated samples were treated with 3AC (5 µM; SHIP1 inhibitor). **(A)** After 24 h, monocyte viability was measured by flow cytometry using annexin V and PI staining. Staurosporine (0.5 µM, ST; non-selective protein kinase inhibitor) was used as a positive control for cell death. **(B)** After 2 h, cells were treated with proteinase K (1.0 mg/ml, 5 min at 37°C) and then washed twice with PBS to remove the viral particles that remained outside of the cell membrane. Samples were then lysed and the viral genome was detected using RT-qPCR (UL123/GAPDH). Statistical significance was measured using one-way ANOVA; **P < 0.005, *P<0.05.

**Table S1.** Antibodies Used in this study.

| Antibody Name | Catalog Number | Note |
| --- | --- | --- |
| Phospho-Akt (Ser473) | Cell Signaling Technology #9271 | Antibodies used for western blot. All the antibodies for western blot was used at 1:1000 concentration other than the antibodies purchased from Santa Cruz, which were used at 1:100 concentration. Rhodamine actin antibody was used at 1:2500 concentration. |
| Phospho-Akt (Thr308) | Cell Signaling Technology #4056 |  |
| Akt (pan) | Cell Signaling Technology #2920 |  |
| SHIP1 | Cell Signaling Technology #2725 |  |
| Phospho-SHIP1 (Tyr1020) | Cell Signaling Technology #3941 |  |
| EGFR | Cell Signaling Technology #4267 |  |
| EGFR | ThermoFisher Scientific #MA5-13070 |  |
| Phospho-EGFR (Tyr1068) | Cell Signaling Technology #3777 |  |
| PDK1 | Cell Signaling Technology #3062 |  |
| Rictor | Cell Signaling Technology #2114 |  |
| Phospho-PRAS40 | Cell Signaling Technology #13175 |  |
| PRAS40 | Cell Signaling Technology #2610 |  |
| Phospho-p70 S6 Kinase (Thr389) | Cell Signaling Technology #9205 |  |
| p70 S6 Kinase | Cell Signaling Technology #9202 |  |
| Phospho-EGFR (Tyr1068) | Abcam #40815 |  |
| mCherry | Abcam #213511 |  |
| Flag | MilliporeSigma #F1804 |  |
| pp28 | Santa Cruz #69749 |  |
| Rhodamine actin | Bio-rad #12004163 |  |
| IE1 | N/A | A generous gift from Tom Shenk (Princeton University). |
| IE2 | N/A |  |
| pUL38 | N/A |  |
| pUL69 | N/A |  |
| pUL98 | N/A |  |
| pp28 | N/A |  |
| gL | N/A | Generated through GenScript custom antibody services and used for western blot. Peptide sequence: KQTRVNLPAHSRYGPQAVDAR |
| CD86 | Biolegend #374214 | Conjugated with BV605 and used for flow cytometry. |
| CD163 | Biolegend #333614 | Conjugated with PE/Cy7 and used for flow cytometry. |

